# Novel RNA Polymerase I and Cyclin Dependent Kinase combination therapy for the treatment of aggressive Acute Myeloid Leukemia

**DOI:** 10.1101/2025.07.21.664618

**Authors:** Nadine Hein, Perlita Poh, Lorena Núñez Villacís, Sarah Sant’Anna Maranhão, Maurits Evers, Jirawas Sornkom, Jason Powell, Priscila Soo, Adria Closa, Jasmina Frawley, Sheren J. Al-Obaidi, Cathryn M. Gould, Piyush Madhamshettiwar, Kaylene J. Simpson, Dani Tutuka, Stuart M Pitson, Luc Furic, Konstantin Panov, Katherine M. Hannan, Amee J. George, Ross D. Hannan, Rita Ferreira

## Abstract

Despite advances in therapy, specific subtypes Acute Myeloid Leukaemia (AML) remains largely incurable. The first-in-class RNA Polymerase I (Pol I) inhibitor, CX-5461, has demonstrated promising activity in both haematological malignancies and solid tumours by selectively inhibiting ribosome biogenesis to induce nucleolar stress, a critical vulnerability in rapidly proliferating cancer cells. CX-5461 has undergone clinical trials for the treatment of both solid and haematological malignancies, and 2^nd^ generation Pol I inhibitor PMR-116 has also entered clinical trials, underscoring the translational potential of drugs that target RNA Polymerase I.

To enhance the therapeutic efficacy of Pol I inhibition and prevent the emergence of resistance, we conducted a cell line-based unbiased screen of FDA-approved drugs to identify compounds that might synergise with CX-5461. This screen revealed that the pan-CDK inhibitors Dinaciclib and Flavopiridol enhance nucleolar stress pathway (NSP) activation and are strong candidates for combinatorial therapy with CX-5461.

Further analysis showed that the combination of CX-5461 and Dinaciclib acts synergistically across a genetically diverse panel of human AML cell lines. This synergy is dependent on an intact NSP, with both agents independently stabilising p53, but with distinct phenotypic outcomes: Dinaciclib induces rapid apoptosis, whereas CX-5461 primarily enforces cell cycle arrest. This functional complementarity results in efficient tumour cell clearance and likely accounts for the delayed onset of therapy resistance. Importantly, combination treatment with CX-5461 and Dinaciclib significantly improved survival in murine models of AML and reduced colony formation in primary human AML samples.

Together these findings provide preclinical evidence for a novel combination treatment strategy that leverages nucleolar stress and cell cycle control to enhance treatment outcomes in AML, paving the way for clinical translation of Pol I-CDK co-targeting therapies.

## Introduction

Given its essential role in sustaining protein synthesis and cell proliferation, rDNA transcription is tightly regulated in normal cells but is frequently dysregulated in cancer, where elevated ribosome production supports malignant growth. This oncogenic dependence on elevated ribosome biogenesis (RiBi) has positioned Pol I transcription as a compelling new therapeutic target^1^. CX-5461 is a first-in-class selective Pol I inhibitor developed to exploit this vulnerability. It acts by preventing the formation of the transcription pre-initiation complex through disruption of the interaction between SL-1 and Pol I at the rDNA promoter^2^. Since its initial discovery, CX-5461 has demonstrated potent anti-tumour activity across a range of pre-clinical cancer models, including haematological malignancies and solid tumours^3–5^. CX-5461 has completed Phase I clinical trials in patients with advanced cancers, where it was shown to be well tolerated, with palmar-plantar erythrodysesthesia reported as the primary adverse event^6,7^. In recognition of its therapeutic promise, CX-5461 received FDA fast-track designation in 2022 for the treatment of homologous recombination deficient breast and ovarian cancers and is undergoing further trial as single agent or in combination (NCT04890613; NCT06606990).

Mechanistically, the on target anti-cancer activity of CX-5461 is primarily driven by acute activation of the nucleolar surveillance pathway (NSP) as a result of disruption to RiBi, rather than delayed effects from ribosome depletion^1^. In the canonical NSP, when Pol I transcription is inhibited, ribosomal proteins such as RPL5 and RPL11 accumulate outside the nucleolus and bind to MDM2, inhibiting its E3 ligase activity and leading to stabilisation of the tumour suppressor protein p53. This triggers downstream responses such as cell cycle arrest, senescence, and/or apoptosis, depending on the genetic context^8^. Notably, NSP activation also occurs through p53-independent “non-canonical” pathways including other ribosomal proteins^9^ and transcriptional regulators such as MYC, E2F, and NF-κB^10^, explaining the heterogeneity of responses observed following Pol I inhibition. In p53 wild-type tumours, CX-5461 typically induces p53-dependent apoptosis, whereas in p53-deficient cells, senescence or cell cycle arrest is often the primary response^5^.

In addition to its effects on Pol I, CX-5461 has been shown to inhibit topoisomerase IIα (Top2A) activity and induce DNA damage^11–13^, highlighting the need to balance efficacy with off-target liabilities. These findings have informed two complementary translational strategies: (i) the development of second-generation Pol I inhibitors with improved selectivity, and (ii) rational design of combination therapies that enhance efficacy at lower doses, thereby reducing off-target effects. Combination therapy represents a particularly attractive strategy in oncology, enabling the simultaneous targeting of parallel or convergent survival pathways and mitigating the risk of therapeutic resistance.

Acute Myeloid Leukemia (AML) is an aggressive form of haematological cancer with increased incidence and mortality in adults over 65 years of age and positive treatment outcome decreases with age^14^. AML is a heterogenous group of diseases and therapeutic options are limited. The current standard of care consists of intensive induction chemotherapy with anthracycline and cytarabine followed by consolidation therapy^15^. However, due to comorbidities associated with age, many patients are ineligible of this treatment. Furthermore, novel treatment options, like the BCL-2 inhibitor Venetoclax, are often ineffective in patients with FLT3-ITD or TP53 mutations, which are associated with resistance and poor prognosis. Therefore, novel approaches that improve treatment efficacy and overcome primary resistance are urgently needed.

Preclinical studies have demonstrated that CX-5461 has activity in AML models^5^; however, strategies to optimise its clinical utility remain underexplored. In this study, we conducted an unbiased screen of FDA-approved compounds to identify agents that synergise with CX-5461 via potentiation of the NSP. This screen identified the cyclin-dependent kinase (CDK) inhibitors Dinaciclib and Flavopiridol as promising candidates. We demonstrate that CX-5461 in combination with Dinaciclib significantly improves overall survival in murine AML models and suppresses colony formation in primary human AML samples. Importantly, this effect is dependent on an intact NSP, supporting nucleolar stress as a tractable therapeutic vulnerability in AML and providing a strong rationale for further clinical development of this combination strategy including with novel second generation inhibitor PMR-116.

## Results

### Co-treatment with CX-5461 and CDK inhibitors exacerbates NSP activation

Inhibition of Pol I transcription by CX-5461 disrupts ribosomal RNA (rRNA) synthesis, thereby activating the NSP. This stress response is marked by stabilisation of the tumour suppressor p53 and induction of its transcriptional programme, resulting in cell cycle arrest, senescence, and/or apoptosis^1^.

To identify compounds that synergise with CX-5461 by further enhancing NSP activity, we established a robust genetic model of nucleolar stress in A549 lung carcinoma cells, a well-recognized system to decipher canonical and non-canonical NSP activation^16^. In this model, knockdown of the ribosomal protein RPS19 via siRNA perturbs RiBi and robustly induces p53 stabilisation, phenocopying RiBi disruption induced p53 accumulation by CX-5461 (Figure 1A-B and quantification in Figure S1A-B). Importantly, this model is well suited for high-throughput screening and allows assessment of NSP activation independently of CX-5461-induced DNA damage.

**Figure 1.**
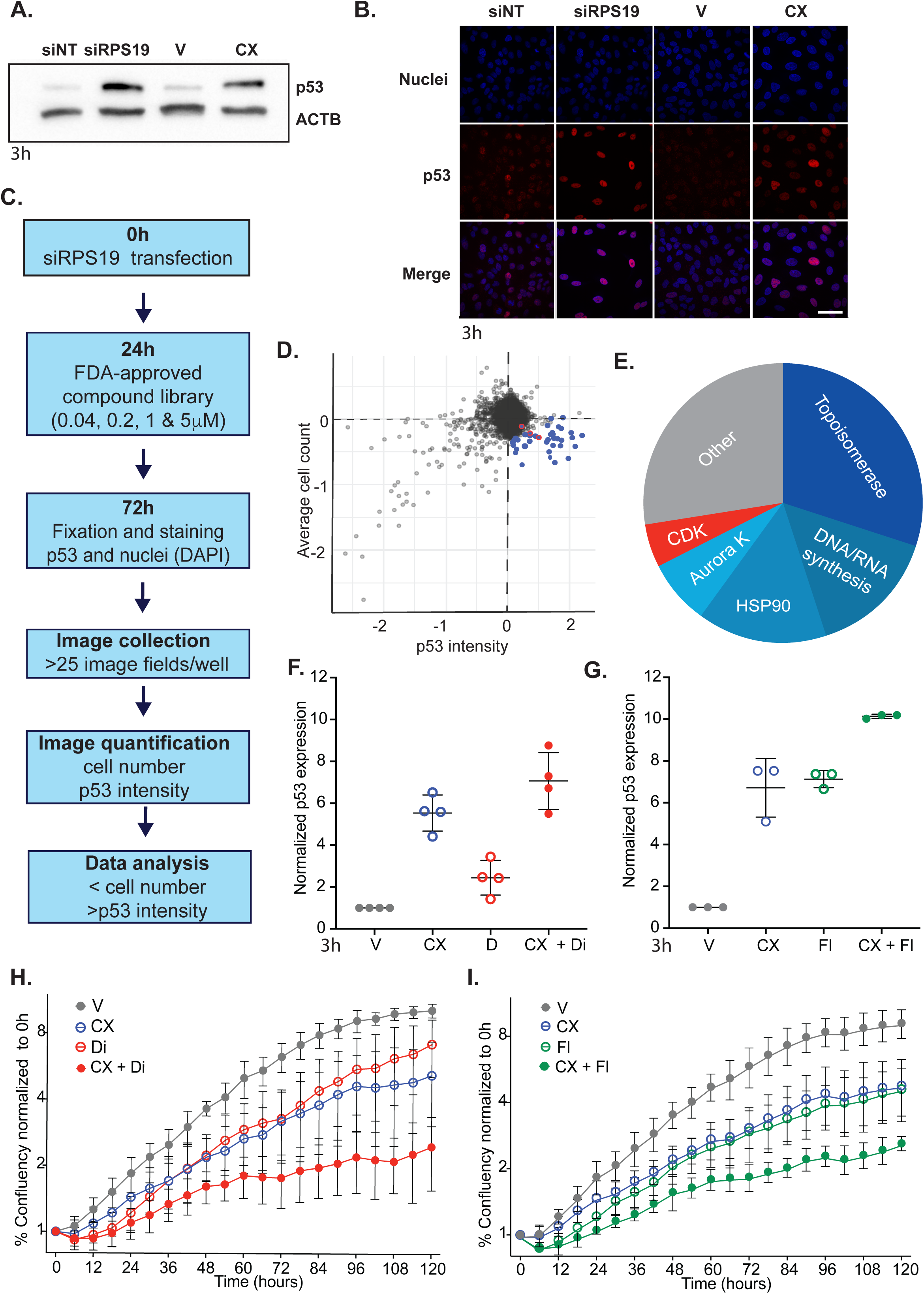
CDK inhibitors Dinaciclib and Flavopiridol are potential combinatorial therapies with CX-5461. **A-B.** p53 expression analysis by western blot (A) and immunofluorescence (B) upon RPS19 KD and CX-5461 treatment. **C.** Experimental design to screen for drugs increasing NSP activation. A549 cells were reverse transfected with Rps19 siRNA for a total of 72 hours. After 24 hours, the media was changed and the compound added for a further 48 hours. The cells were then fixed and stained with a nuclear marker (DAPI) to quantify the number of cells, and p53 antibody for p53 intensity, using immunofluorescence analysis. Image analyses of these microplates was then performed on the Cellomics ArrayScan VTi with two channels: channel 1 for DAPI and channel 2 for p53 intensity. For each well, 25 fields were captured and analysed to collect information on cell number and intensity of p53 staining. **D.** Distribution of Mahalanobis distance of each library compound across all drug concentrations. Top ranking compounds causing increase of p53 intensity and reduction in cell numbers are highlighted in blue. Data is normalised to A549 RPS19 KD treated with DMSO. **E.** Categories of top 40 ranked compounds inducing stability of p53 and decreasing numbers across all concentrations. CDK: cyclin dependent kinase; HSP: Heat shock protein; **F-G.** Treatment of A549 cells for 3h with TIC50 concentrations of CX-5461 (CX; 400nM) and Dinaciclib (Di; 20nM) (F) or CX and Flavopiridol (Fl; 300nM) (G) has an additive effect when compared to single drug treatments. display mean ± standard deviation of at least 3 biological replicates. **H-I** Percentage confluence over time, normalised to time 0, of A549 cells treated with (H) 175nM CX + 15nM Di and (G) 150nM CX + 200nM Fl. Graphs represent mean +/− SEM of 3 independent experiments. Significance was calculated using ordinary one-way ANOVA with Bonferroni’s multiple comparison test. * p<0.5, ** p<0.05, ***p<0.005.

Using this system, we screen a library of 4169 FDA-approved drugs across four concentrations (ranging from 40nM to 5μM). High-content imaging was utilized to quantify changes in p53 abundance and cell number, estimated via nuclear staining (Figure 1C). Drugs were scored according to their potential for NSP activation, as evidenced by enhanced p53 stabilisation, and further reductions in cell viability beyond that observed in RPS19 knockdown alone (Figure 1D; Figure S1C).

The top 40 compounds that consistently enhanced NSP activation and decreased cell number across all tested concentrations were classified according to their known molecular targets (Figure 1E, Table 1). The largest group of hits targeted topoisomerases (13 compounds; 32.5%), followed by inhibitors of RNA/DNA synthesis (6 compounds; 15%), HSP90 inhibitors (6 compounds; 15%), cyclin-dependent kinase (CDK) inhibitors (3 compounds; 7.5%), and Aurora kinase inhibitors (2 compounds; 5%). The remaining compounds (27.5%) were single agents targeting other pathways.

**Table 1.** Top 40 ranking compounds inducing p53 stabilization of and decrease in cell number.

|  | Drug Name | Target |
| --- | --- | --- |
| 1 | 17-DMAG | HSP90 |
| 2 | Camptothecine_(S,+) | Topoisomerase I |
| 3 | Ganetespib (STA-9090) | HSP90 |
| 4 | Mitoxantrone dihydrochloride | DNA topoisomerase II |
| 5 | Mitoxantrone | DNA synthesis |
| 6 | Camptothecin | DNA topoisomerase II |
| 7 | Mitoxantrone | Topoisomerase |
| 8 | AT13387 | HSP90 |
| 9 | Adriamycin | Topoisomerase |
| 10 | NVP-AUY922 | HSP90 |
| 11 | Doxorubicin hydrochloride | DNA intercalant |
| 12 | (S)-(+)-Camptothecin | DNA topoisomerase I |
| 13 | Camptothecine | Topoisomerase |
| 14 | Actinomycin D | RNA polymerase |
| 15 | Clofarabine | ribonucleotide reductase |
| 16 | 17-AAG(Geldanamycin) | HSP90 |
| 17 | BIIB021 | HSP90 |
| 18 | RITA | p53-MDM2 interaction |
| 19 | Daunorubicin hydrochloride | DNA intercalating |
| 20 | Cladribine | adenosine deaminase |
| 21 | Topotecan hydrochloride hydrate | topoisomerase I |
| 22 | ABT-418 hydrochloride | Nicotinic acetylcholine receptor |
| 23 | Doxorubicin hydrochloride | DNA topoisomerase II |
| 24 | ABT-869 | RTK, VEGFR, PDGFR, FGFR |
| 25 | **Dinaciclib** | **CDK** |
| 26 | **Flavopiridol** (Alvocidib) | **CDK** |
| 27 | PHA-680632 | Aurora Kinase |
| 28 | Nitrofurantoin | Bacterial DNA damage |
| 29 | Fludarabine | DNA synthesis |
| 30 | cis-4-Aminocrotonic_acid | GABA-C receptor |
| 31 | BIO | GSK-3 |
| 32 | Daunorubicin hydrochloride | RNA synthesis |
| 33 | PHA-739358 (Danusertib) | Aurora Kinase, Bcr-Abl |
| 34 | Hydroxytacrine maleate | Acetylcholine esterase |
| 35 | **Flavopiridol** (HMR-1275; Alvocidib; L868275) | **CDK** |
| 36 | Fananserin | 5-HT<sub>2A</sub>; D<sub>4</sub> |
| 37 | XL147_(SAR245408) | PI3K |
| 38 | A61603 hydrobromide | alpha1A |
| 39 | SN38 | DNA topoisomerase I |
| 40 | Docetaxel | Microtubule |

We selected the pan-CDK inhibitors (CDKi) for further mechanistic analysis as previous work from our group has demonstrated synergy between topoisomerase inhibitors (including Topotecan) and CX-5461 in ovarian cancer models^17^. Similarly, the ability of RNA and DNA synthesis inhibitors to trigger nucleolar stress and activate the NSP is already well established^1,16,18^. Although several HSP90 and Aurora kinase inhibitors have progressed into clinical development, their therapeutic potential has been limited by dose-limiting toxicity^19,20^.

Two CDK inhibitors emerged from the screen, Dinaciclib and Flavopiridol, both of which have advanced to Phase II clinical trials for the treatment of haematologic malignancies^21^. These compounds display preferential inhibition of CDK9, a kinase essential for transcriptional elongation of RNA polymerase II (Pol II). CDK9 phosphorylates serine 2 on the C-terminal domain of Pol II (RPB1), a modification required for productive transcription elongation^22,23^. CDK9 inhibitors have been shown to have efficacy in the treatment of AML through the suppression Pol II-driven expression of short-lived survival proteins, such as MCL-1 and MYC^24^. We therefore hypothesised that concurrent inhibition of Pol I (via CX-5461) and Pol II (via CDK9 inhibition) would yield a potent combinatorial effect, broadening the therapeutic window and potentially delaying the onset of resistance.

Having confirmed that both Dinaciclib and Flavopiridol further enhance p53 stabilisation in the RPS19 knockdown A549 cell system used for our initial screen (Figure S1D), we next evaluated their combinatorial potential with CX-5461. To do so, we assessed molecular markers of drug activity in A549 cells treated with single agents or in combination. The 50% target inhibitory concentration (TIC50) of CX-5461, Dinaciclib and Flavopiridol targets, pre-rRNA and CDK9 phosphorylation of RPB1, respectively, were determined in A549 (Figure S1E-F). A549 cells were treated for 3h with vehicle or TIC50 concentrations of CX-5461 (750nM), Dinaciclib (20nM) or Flavopiridol (300nM) alone or in combination. In line with our hypothesis, co-treatment with CX-5461 and either Dinaciclib or Flavopiridol resulted in an additive increase in p53 stabilisation compared to single-agent treatments (Bliss synergy score<10; Fig 1F-G), consistent with enhanced activation of the NSP.

To evaluate the impact of CX-5461 and CDK inhibitor co-treatment on cell proliferation, A549 cells were treated with vehicle, CX-5461 (150-200nM), Dinaciclib (10-20nM), or Flavopiridol (100-200nM), either alone or in combination. These drug concentrations were selected based on pre-determined growth inhibitory concentrations (GIC₅₀) for each compound (CX-5461: 175nM; Dinaciclib: 15nM; Flavopiridol: 200nM; Figure S1H-I). Live-cell imaging was performed every 6 hours until the vehicle-treated control reached 95% confluency. To assess drug interaction, Bliss synergy scores were calculated using SynergyFinder. The strongest synergistic inhibition of cell proliferation was observed with the combination of 175nM CX-5461 and 15nM Dinaciclib (Bliss score=25.0; Figure 1H), followed by 175nM CX-5461 and 100nM Flavopiridol (Bliss score=12.4; Figure 1I), with both Bliss scores being >10 indicating a synergistic rather than merely additive effect of these combinations.

### Synergistic anti-cancer effect of CX-5461 - Dinaciclib combination treatment in multiple AML subtypes

Given that Dinaciclib exhibits greater selectivity for CDK1, CDK2, CDK5, and CDK9, (IC90 in the low nM range) and demonstrates a more favourable toxicity profile compared to the broader-spectrum CDK inhibitor Flavopiridol^25^, we prioritised further cell line evaluation of the CX-5461 and Dinaciclib combination.

We compiled a panel of AML cell lines, encompassing diverse driver mutations and including both TP53 wild-type and TP53-null backgrounds, to further investigate the therapeutic potential of CX-5461-Dinaciclib combination therapy. Cells were treated for 72 hours with vehicle, or 0.1×, 1×, and 10× the IC₅₀ of CX-5461 and Dinaciclib (determined by^26^ and^24^, respectively) alone or in combination, and cell viability was assessed by PI exclusion using flow cytometry (Figure 2A; Figure S2A). Chou-Talalay combination index (CI) was calculated for effect levels between 50 and 90% (fraction affected 0.5-0.9) and a conbinatorial effect was considered synergistic if a single CI was lower than 1. Synergy was identified in 5 out of 6 AML cell lines independently of p53 status (Figure 2B). Synergy was observed in MV4;11 (CI=0.64±0.135; p53 WT), NOMO-1 (CI=0.61±0.067;p53 mutant), MOLM-13 (CI=0.68±0.383;p53 WT), KG1 (CI=0.71±0.350; p53 null) and SHI-1 (CI=0.70±0.626) but not in OCI-AML3 (CI=0.994±0.452). MV4;11 cells were selected for further mechanistic analysis as they exhibited consistently low CI at all effect levels indicating strong synergy between CX-5461 and Dinaciclib and have a functional NSP (p53 WT). Collectively, these data indicate that the combination of CX-5461 and Dinaciclib represents a promising therapeutic strategy across genetically diverse AML subtypes, irrespective of driver mutation or p53 status.

**Figure 2.**
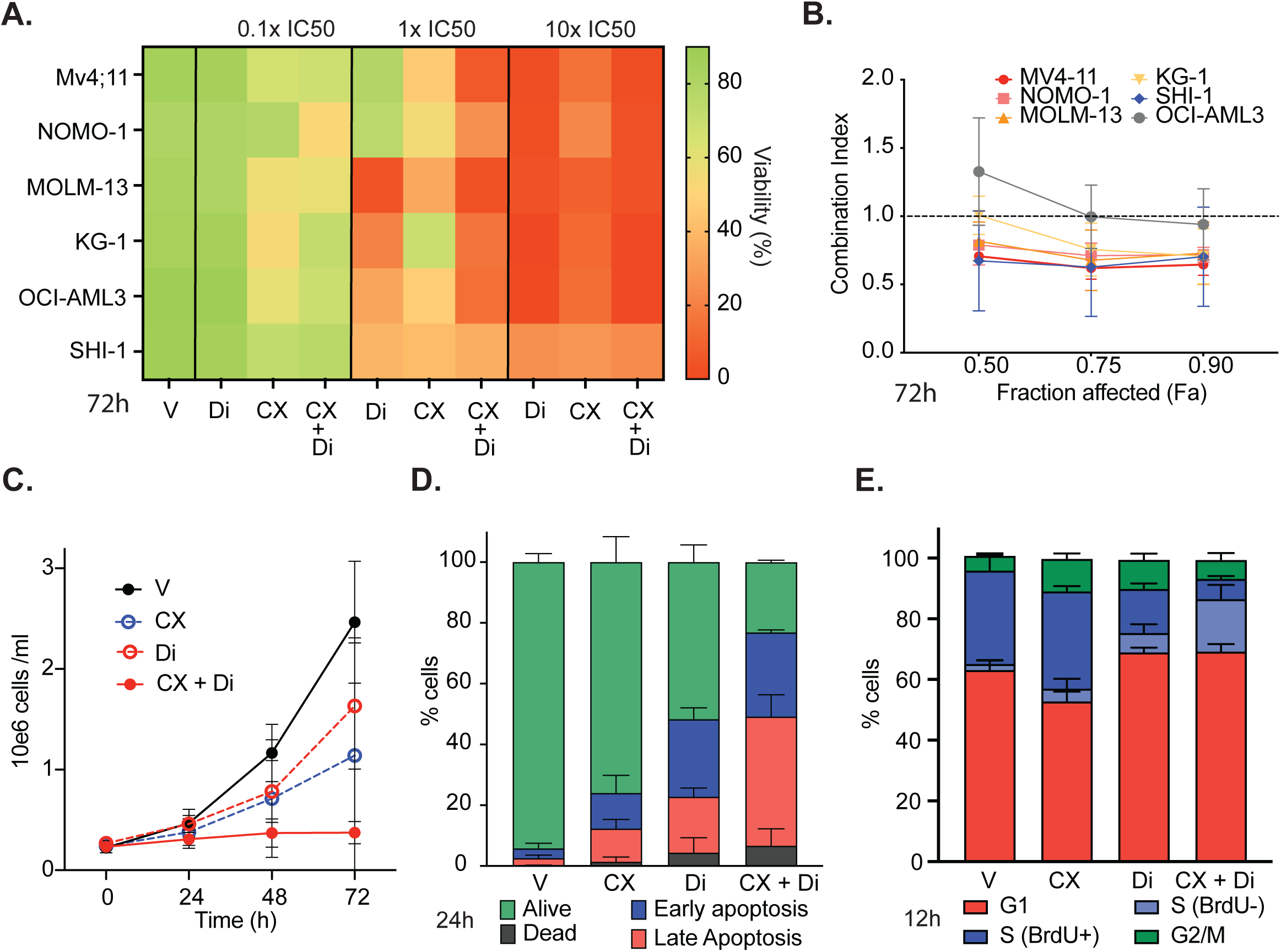
CX-5461 and Dinaciclib combinatorial therapy is effective in a variety of human AML cell lines. **A.** Matrix showing percentage viability, determined by Propidium Iodine exclusion, of a panel of 6 cell lines (Mv4;11, Nomo-1, MOLM-13, KG1. OCMI-AML3 and SHI-1) treated for 48 hours with 0.1X, 1X and 10X GIC50 concentrations of Cx-5461 and Dinaciclib. **B.** Combination index (CI) analysis of the CX-5461 and Dinaciclib treated AML cell line panel. **C.** Growth curves of Mv4;11 cells treated with TIC50 concentrations of CX-5461, Dinaciclib or combination of both for 3 hours. **D.** Apoptosis analysis of MV4;11 treated CX-5461, Dinaciclib or combination of both for 24 hours. **E.** Cell cycle analysis of MV4;11 treated CX-5461, Dinaciclib or combination of both for 12 hours. All graphs display mean ± standard deviation of at least 3 biological replicates. Significance was calculated using ordinary one-way ANOVA with Bonferroni’s multiple comparison test. * p<0.5, ** p<0.05, ***p<0.005.

### CX-5461 and Dinaciclib combinatorial treatment induces proliferation arrest and cell death

MV4;11 is an aggressive AML cell line characterised by the presence of the MLL-AF4 fusion and an internal tandem duplication (ITD) in FLT3, genetic features associated with poor prognosis and high proliferative potential. To assess mechanistically how CX-5461 and Dinaciclib combination treatment altered proliferation capacity, MV4;11 cells were treated for 3 hours with previously established TIC₅₀ of CX-5461 (300nM; Figure S2B) and Dinaciclib (10nM; Figure S2C-D), either as monotherapies or in combination. Under these conditions TIC₅₀ concentrations of the single agents caused a moderate (CX-5461-56%; Dinaciclib - 36%) reduction in cell proliferation; however, co-treatment resulted in a complete proliferation arrest (Figure 2C).

The induction of apoptosis was quantified at 3, 6, and 24 hours post-treatment using Annexin V and 7AAD staining assessed by flow cytometry. No significant increase in apoptotic cells was observed at 3- and 6-hours post-treatment (Figure S2E), however, by 24 hours, a significant increase in the proportion of cells undergoing early (Annexin V^+^, 7AAD^−^) and late apoptosis (Annexin V^+^, 7AAD^+^) was observed with both CX-5461- (10.9±2.98%;11.7±5.81%) and Dinaciclib-treated (18.3±2.90%; 25.5%±3.75%) cells compared to vehicle treated cells (2.4%±0.98%; 3.2%±1.70%). A synergistic increase in late apoptotic cells (42.5±7.21%; Excess over Bliss CI=0.299 (0.170-0.429)) was observed with the combination treatment (Figure 2D).

To assess the effect of treatment on cell cycle progression, cells were analysed 12 hours post-treatment, prior to the onset of detectable cell death. Cells were pulsed with BrdU for 30 minutes, and analysed by flow cytometry. As previously reported, CX-5461 treatment delayed S-phase progression and led to an accumulation of cells in the G2/M phase (10.7±0.87% vs 4.85±1.84% with vehicle). Dinaciclib treatment significantly reduced the S-phase population (20.9±4.69% vs 32.8±6.39% with vehicle) and induced a modest accumulation in G1 phase (68.9±1.56% vs 63.0±3.35% with vehicle). Interestingly, the combination treatment also reduced the overall number of S-phase cells (23.9±5.59% vs 32.8±6.39% with vehicle), but this was due to S-phase arrest rather than delayed progression, as indicated by an increased proportion of BrdU-negative S-phase cells (17.3±4.65% vs 2.1±0.87% with vehicle). A modest increase in the G1 population was also observed following combination treatment (69.2±2.41% vs 63.0±3.35% with vehicle; Figure 2E). Cell cycle analysis at 24 hours post-treatment yielded similar results, although a reduction in viable cells due to extensive apoptosis limited interpretation (Figure S2F).

Collectively, these data support the conclusion that the in vitro combinatorial effect of CX-5461 and Dinaciclib is driven primarily by enhanced induction of apoptosis, coupled with cell cycle arrest.

### Distinct but Convergent On-Target Mechanisms Drive Synergistic Anti-Leukemic Activity of CX-5461 and Dinaciclib

To explore the mechanistic basis for the efficacy of CX-5461 and Dinaciclib combination therapy, we first examined whether each compound engages its known molecular targets and downstream effectors.

As expected, CX-5461 monotherapy significantly suppressed 47S pre-rRNA expression, consistent with its inhibition of RNA Polymerase I transcription. Notably, this suppression was further enhanced by co-treatment with Dinaciclib, indicating an additive effect on rRNA synthesis. While Dinaciclib alone reduced pre-rRNA levels in four out of six assays, variability in the response rendered this effect statistically non-significant (Figure 3A). Although CDK9 inhibition has been reported to affect rRNA synthesis at higher doses^27^, such an effect was not anticipated under the dosing conditions used here (Figure S2D). Furthermore, we found that neither Dinaciclib nor Flavopiridol impaired the catalytic activity of Pol I or its ability to elongate transcripts, across a wide range of concentrations, as evidenced by the promoter-independent in vitro transcription assay (non-specific assay, NSA; Figure S3A). Similarly, no significant inhibitory effect was observed in the promoter-driven run-off transcription assays (Promoter specific assay, PSA), indicating that the compounds did not interfere with the assembly of the Pol I pre-initiation complex or the initiation of transcription (Figure S3B).

**Figure 3.**
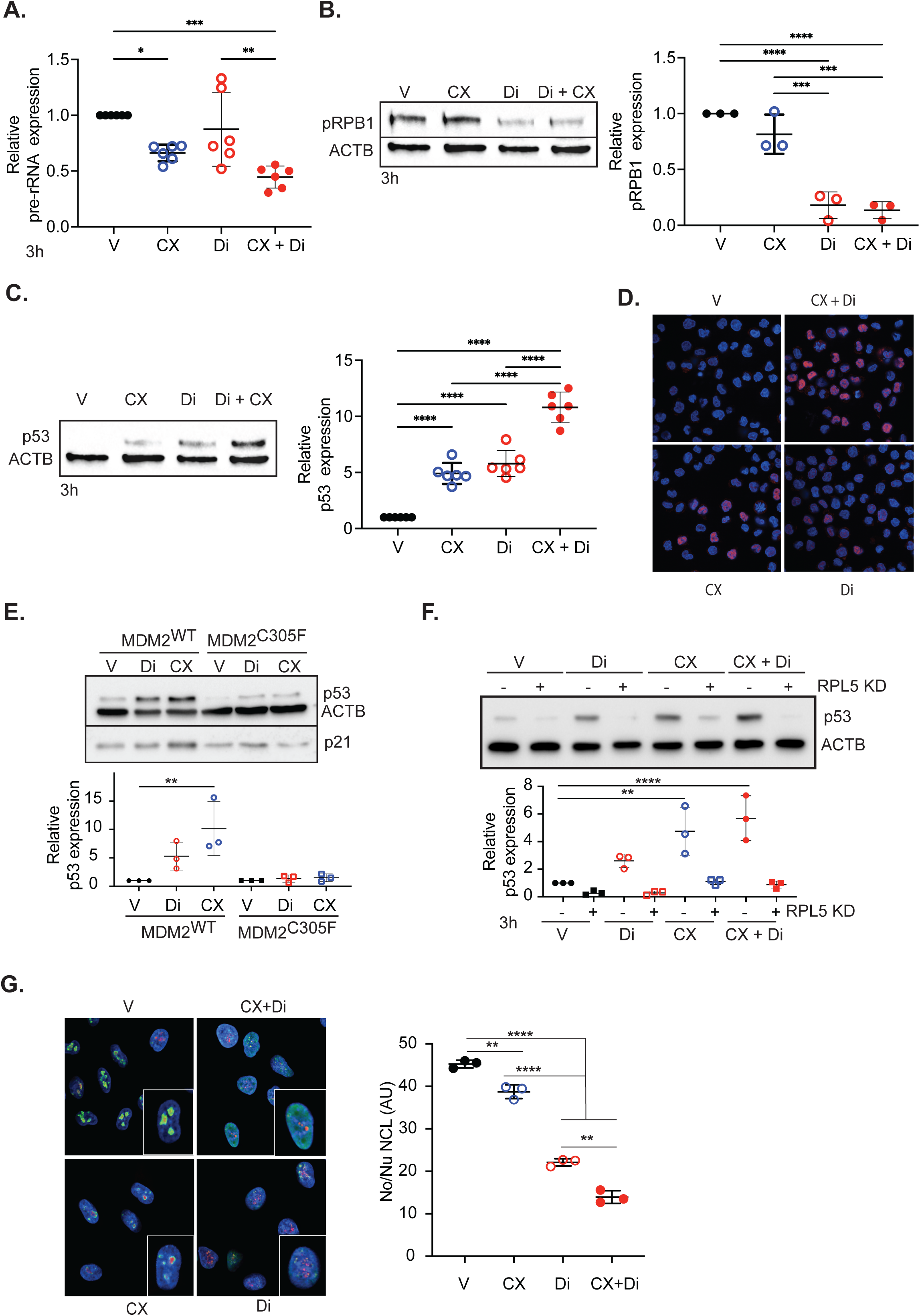
CX-5461 and Dinaciclib combinatorial effect is driven by the activation of the canonical NSP. Mv4;11 cells were treated for 3h with vehicle and TIC50 concentrations of CX-5461 (300nM), Dinaciclib (10nM) or combination of both drugs **A.** Pre-rRNA expression levels, normalized to housekeeping gene B2M and Vehicle and **B.** Representative immunoblot for Serine 2 phospho-RPB1 and ACTB protein levels (left) and protein levels normalized to loading control ACTB and vehicle (right). **C.** Representative immunoblot for p53 and ACTB protein levels (left) and protein levels normalized to ACTB and vehicle (right). **D.** Representative images of p53 (red) and DAPI (cyan) immunofluorescence. **E.** Representative immunoblot of p53 and p21 protein (top) and quantification of p53 levels normalized to BACT and vehicle (bottom) in wild type and MDM2^C305F^ mutant MEFS upon 3h treatment with TIC50 concentration of CX-5461 or Dinaciclib. **F.** Representative immunoblot of p53 protein levels protein (top) and quantification of p53 levels normalized to BACT and vehicle (bottom) in A549 with and without knockdown of RPL5. **G**. Representative images of Fibrillarin (red), Nucleolin (green) and DAPI (cyan) immunofluorescence (left) and quantification of Nucleolin dispersal (right) in A549 cells treated for 3h with GIC50 concentrations of CX-5461 (175nM) and Dinacicib (15nM), or combination of both drugs. All graphs display mean ± standard deviation of at least 3 biological replicates. Significance was calculated using ordinary one-way ANOVA with Bonferroni’s multiple comparison test. * p<0.5, ** p<0.05, ***p<0.005.

Phosphorylation of RPB1 at Ser2, catalyzed by CDK9 to promote transcriptional elongation by RNA Polymerase II, serves as a surrogate marker for on-target CDK9 inhibition. Accordingly, Dinaciclib treatment led to a marked reduction in Ser2 phosphorylation, confirming effective target engagement (Figure 3B). A comparable reduction was observed in the combination-treated cells, while CX-5461 had no effect on RPB1 phosphorylation, as expected.

In parallel, we assessed activation of the canonical NSP, a p53-dependent stress pathway triggered by impaired ribosomal RNA synthesis. CX-5461 monotherapy induced robust (∼5x over vehicle) p53 stabilization, consistent with its Pol I inhibitory activity. Dinaciclib also induced p53 accumulation (∼5x over vehicle), consistent with prior reports of CDK9 inhibition affecting p53 regulators. Co-treatment resulted in a synergistic increase in p53 protein levels (∼11x over vehicle; Figure 3C), corroborated by immunofluorescence staining (Figure 3D; quantification in Figure S3C).

To determine the extent by which p53 response was NSP-dependent in response to both drugs, we employed mouse embryonic fibroblasts (MEFs) from the Mdm2^C305F^ mutant mouse strain^28^, in which the canonical NSP is genetically disrupted due to impaired binding of the 5S ribonucleoprotein (5S-RNP) complex to MDM2. In these cells, treatment with CX-5461 or Dinaciclib, alone or in combination, did not increase p53 abundance, whereas wild-type MEFs exhibited a 5- to 10-fold increase with the combination (Figure 3E). These findings were independently validated by siRNA-mediated knockdown of RPL5 or RPL11 in A549 cells, which similarly abolished p53 stabilisation in all treatment conditions (Figure 4F; Figure S3D). These results demonstrate that p53 activation by both agents is mediated via the canonical NSP and not through generalized cellular stress, highlighting Dinaciclib’s unexpected engagement of this pathway.

**Figure 4.**
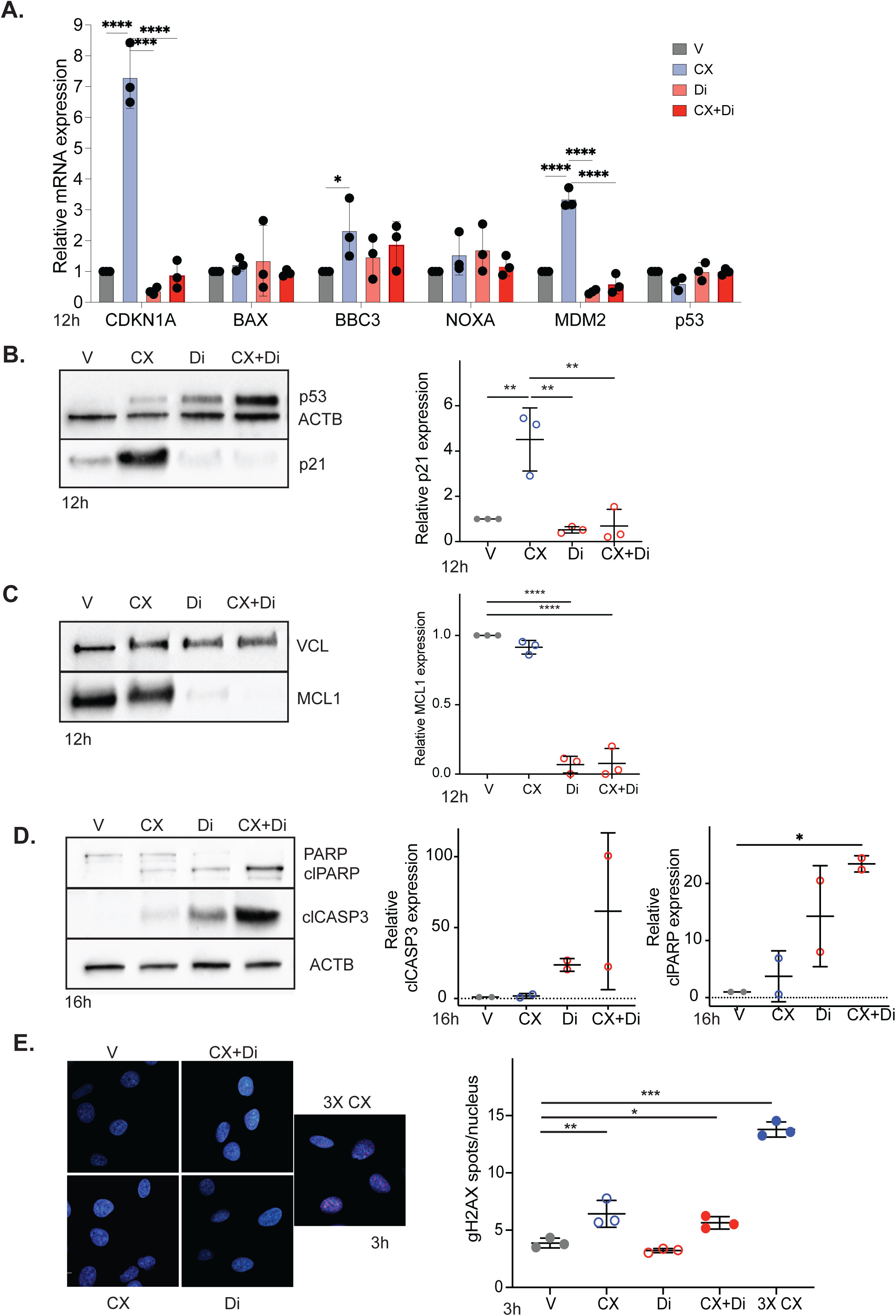
Distinct mechanisms of CX-5461 and Dinaciclib drive their synergistic effect. **A.** Relative mRNA expression analysis of p53 target genes, normalized to housekeeping gene B2M and vehicle, in Mv4;11 cells treated for 3h with vehicle and TIC50 concentrations of CX-5461 (300nM), Dinaciclib (10nM) or combination of both drugs **B.** Representative immunoblot of p53 and p21 protein (left) and quantification of CDKN1 (p21) levels normalized to BACT and vehicle (right). **C.** Representative immunoblot of MCL1 and Vinculin (VCL) proteins (left) and quantification of p21 levels normalized to VCL and vehicle (right). **D.** Representative immunoblot of cleaved Caspase 3 (clCASP3) and full length and cleaved PARP (clPARP) proteins (left), quantification of clCASP3 (centre) and clPARP levels (right) normalized to VCL and vehicle. **E.** Representative images of γH2AX (red) and DAPI (cyan) immunofluorescence treated with C-5461 (175nM) and Dinacicib (15nM) alone, or in combination (left) and quantification of average number of γH2AX foci per nucleus (right) in A549 cells treated for 3h with GIC50 concentrations of CX-5461. All graphs display mean ± standard deviation of at least 3 biological replicates. Significance was calculated using ordinary one-way ANOVA with Bonferroni’s multiple comparison test. * p<0.5, ** p<0.05, ***p<0.005.

Supporting this, nucleolar structure was disrupted 12 hours post-treatment with both agents. Dinaciclib led to dispersal of Nucleolin, consistent with nucleolar disassembly, while CX-5461 caused nucleolar enlargement and peripheral Nucleolin accumulation. Co-treatment led to more extensive nucleolar disruption, quantitatively confirmed by measuring Nucleolin redistribution relative to Fibrillarin-positive regions (Figure 3E). Together, these findings indicate that both drugs converge on NSP activation via distinct upstream mechanisms, contributing to the synergistic stabilization of p53.

To further dissect downstream consequences, we examined key transcriptional and apoptotic targets. In MV4;11 cells treated for 3 hours, CX-5461 alone increased mRNA levels of p53 targets CDKN1A (p21) and MDM2, while no significant changes were observed for pro-apoptotic genes such as BAX, BBC3 (PUMA), or NOXA (Figure 4A), likely due to the early time point and absence of apoptosis (Figure S2F). These transcriptional changes were mirrored at the protein level at both 3h (Figure S4A) and 12 hours (Figure 4B) post-treatment, with ∼5X p21 upregulation restricted to CX-5461-treated cells, suggesting that its primary effect, under this conditions (concentration and time of treatment) is to induce cell cycle arrest or senescence.

In contrast, Dinaciclib downregulated MCL1, a short-lived anti-apoptotic transcript dependent on Pol II elongation. MCL1 mRNA and protein levels were reduced at 6 and 12 hours, respectively, in both Dinaciclib and combination-treated cells (Figure S4B and Figure 4C).

Cleaved Caspase 3 and cleaved PARP proteins, indicators of active apoptosis, were detected at 16 hours post treatment (Figure 4D). CX-5461 treatment led to low levels of cleaved proteins while Dinaciclib treatment led to >20X increase in the levels of detectable cleaved proteins compared to vehicle treated cells. Combinatorial treatment led to a further increase in detectable cleaved proteins (∼60X clCASP3 and ∼27X for clPARP over vehicle) consistent with active Caspase 3 mediated apoptosis.

Collectively, these data reveal that CX-5461 and Dinaciclib activate distinct downstream programs: CX-5461 induces the canonical p53–p21 axis leading to cell cycle arrest, while Dinaciclib suppresses pro-survival transcriptional programs such as MCL1 to trigger apoptosis. Despite their distinct primary mechanisms, Pol I–NSP–p53 versus Pol II–CDK9–MCL1, both agents also converge on the nucleolar stress response, resulting in synergistic tumour cell death. These findings provide a mechanistic rationale for co-targeting Pol I and CDK9 as a therapeutic strategy in AML and other cancers with intact NSP function.

### Reduced CX-5461-Associated DNA Damage with Combination Therapy

Due to the synergistic effect between CX-5461 and Dinaciclib, a lower concentration of each drug can be used, thereby reducing off-target or concentration-dependent side effects. Since CX-5461 is known to induce DNA damage in a concentration-dependent manner, we investigated whether combination treatment reduced the induction of double-strand breaks. A549 cells were treated with CX-5461 (150nM) and Dinaciclib (15nM) for 3 hours, concentrations known to cause a synergistic inhibitory effect on cell proliferation, and the number of DNA double-strand break foci was quantified by immunofluorescence staining of γH2AX. As expected, the average number of γH2AX foci increased upon treatment with CX-5461 alone, although the magnitude was substantially lower than that induced by a higher CX-5461 dose (3× CX) required for effective single-agent treatment (Figure 4E). Treatment with Dinaciclib alone did not induce γH2AX foci, and combinatorial treatment did not further increase the number of foci beyond that seen with CX-5461 alone. These results indicate that combination therapy mitigates CX-5461-induced DNA damage by enabling therapeutic efficacy at reduced dosing.

Together, these complementary mechanisms not only enhance anti-leukemic efficacy but also minimise genotoxic stress, supporting the clinical potential of CX-5461 and Dinaciclib as a well-tolerated and effective therapeutic combination for high-risk AML.

### Combinatorial Inhibition of Pol I and CDK9 Prolongs Survival in Genetically Distinct in vivo Murine AML Models

To evaluate the *in vivo* therapeutic potential of combining CX-5461 with Dinaciclib, which has been trialled individually or in combination with other agents in haematological malignancies^24,29^ we employed a well-established, aggressive murine model of AML. This model involves transplantation of syngeneic recipient mice with AML cells expressing a GFP-tagged MLL-ENL fusion protein and luciferase-tagged oncogenic *Nras*, recapitulating features of high-risk disease^30^. Leukaemia engraftment was detectable at day 7 post-transplantation. On day 8, animals were randomised into treatment cohorts (n=10 per group) and administered either vehicle, single agents (CX-5461 or Dinaciclib or in combination. Doses were selected based on prior tolerability and efficacy data: CX-5461 (30 mg/kg) and Dinaciclib (25 mg/kg). The concentration of CX-5461 is lower that the maximum tolerated dose of 40mg/kg used in prior monotherapy studies^26^.Mice were monitored weekly for disease progression using bioluminescent imaging until ethical endpoints were reached (Figure 5A and Figure S5A).

**Figure 5.**
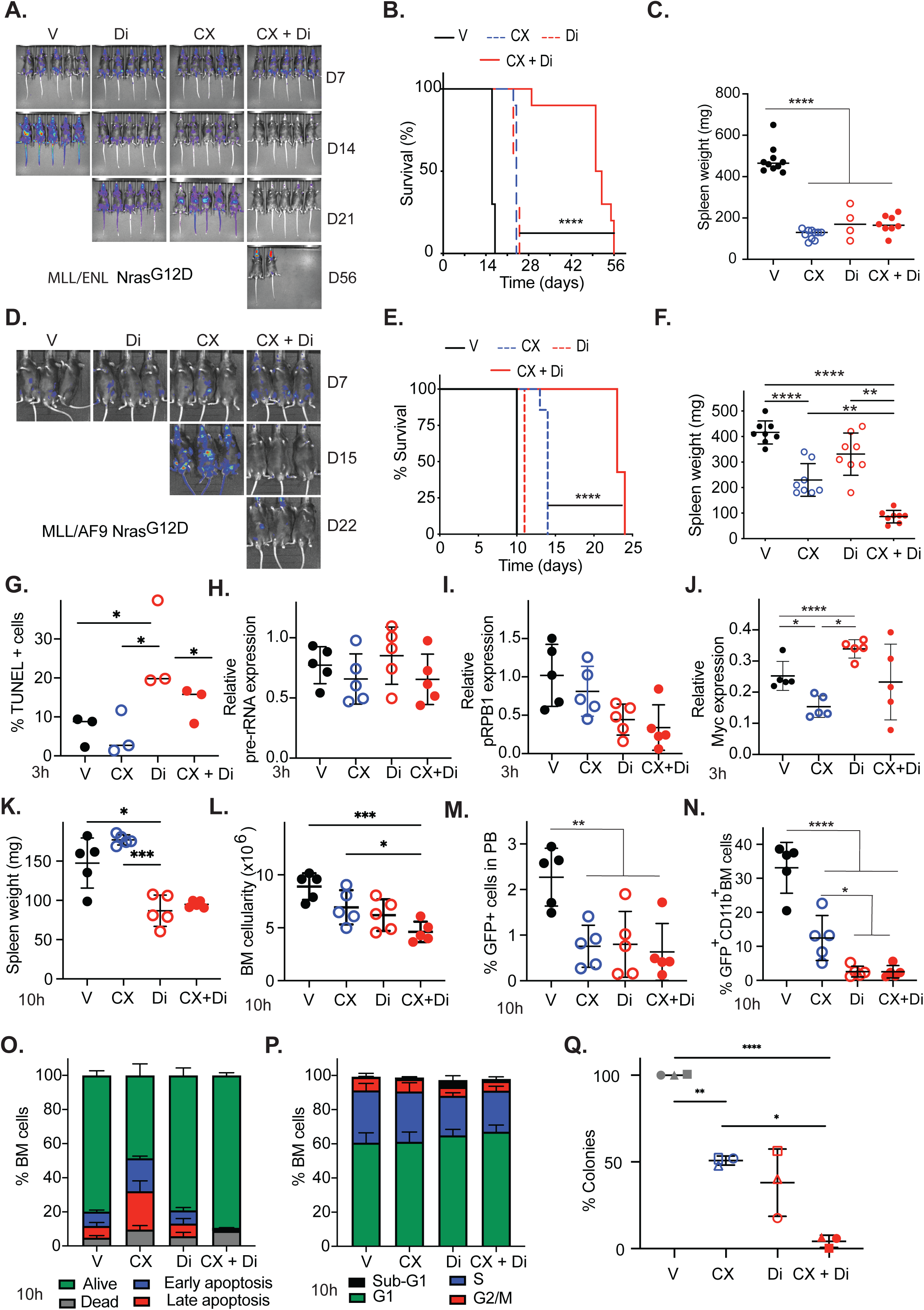
Combinatorial treatment with CX-5461 and Dinaciclib or Flavopiridol is efficacious in AML models. **A.** Representative images of bioluminescence imaging used to monitor disease progression of MLL-ENL Nras AML model treated with vehicle, CX-5461, Dinaciclib or combination therapy **B**. Kaplan-Meier plot showing overall survival of MLL-ENL Nras AML model treated with vehicle, CX-5461, Dinaciclib or combination therapy. Data for the survival study was analysed by log-rank (Mantel-Cox) test with Benjamini and Hochberg correction, n = 10 mice/group. **** p < 0.0001, *** p < 0.001. **C.** Spleen weight measured at ethical endpoint after treatment with either single agent or combination treatment. Data was analysed by ordinary one-way ANOVA multiple comparison test with the Original FDR method of Benjamini and Hochberg, Significance **** p < 0.0001, *** p < 0.001, ** p < 0.01, * p < 0.05. **D.** Representative images of bioluminescence imaging used to monitor disease progression **E**. Kaplan-Meier plot showing overall survival of MLL-AF9 Nras^G12D^ AML model treated with vehicle and **F**. Spleen weights of CX-5461 and Dinaciclib monotherapy, and combination therapy. **G.** Percentage of TUNEL positive spleen cells **H.** Spleen pre-rRNA **I.** Serine 2 phosphorylated RBP1 protein and **J.** MYC protein expression in oMLL-ENL Nras AML model treated for 3h with vehicle, CX-5461, Dinaciclib or combination therapy. **K.** Spleen weight **L.** bone marrow cell number **M.** percentage of tumour (GFP^+^) cells in peripheral blood and **N.** percentage of tumour (GFP^+^CD11B^+^) cells in bone marrow **O.** apoptosis and **P.** cell cycle analysis of MLL-ENL Nras AML model treated for 10h with vehicle, CX-5461, Dinaciclib monotherapy or combination therapy. **Q**. Percentage of patient primary colonies upon treatment with CX-5461, Dinaciclib and combination therapy, normalised to vehicle. All graphs display mean ± standard deviation of at least 3 biological replicates. Significance was calculated using ordinary one-way ANOVA with Bonferroni’s multiple comparison test. * p<0.5, ** p<0.05, ***p<0.005.

Combination treatment with CX-5461 and Dinaciclib resulted in a marked survival benefit compared to vehicle control (p < 0.0001), as well as to single-agent treatment arms: CX-5461 (p < 0.0001) and Dinaciclib (p < 0.0001). Median survival was significantly extended to 32 days when CX-5461 and Dinaciclib were given in combination compared to 24 and 23 days for CX-5461 and Dinaciclib, respectively (Figure 5B). Ethical endpoints were predominantly reached due to focal residual disease in the cranial region, which caused breathing difficulties. Nonetheless, spleen weights at endpoint were significantly reduced in all treatment groups relative to vehicle controls, confirming a reduction in systemic leukemic burden (Figure 5C).

Simmilarly, median increased survival was observed in cohorts treated with Flavopiridol alone (16 day p < 0.001) that was further extended when in combination with CX-5461 (24 days), although at a reduced extent (Figure S5A-C).

To assess the generalisability of the therapeutic effect, we extended our studies to a second aggressive AML model driven by MLL-AF9 and *NRas*^G12D^ mutations. In this model, co-treatment with CX-5461 and Dinaciclib again produced a significant increase in overall survival relative to monotherapy or vehicle-treated controls (Figure 5D). Dinaciclib therapy alone show little survival benefict compared to vehicle treated mice (11 vs 10 days, respectively), while CX-5461 therapy increased survival to 15 days. Notably, mice treated with combinatorial therapy survive to 24 days (Figure 5E) and displayed significantly reduced spleen weights (Figure 5F). A reduction in circulating tumour cells was also observed in mice treated with CX-5461 and combination but not in mice treated with Dinaciclib alone (Figure S5D). Analysis of bone marrow (Figures S5E)and spleen (Figures S5F) show a moderate (∼50%) and variable (not statistically significant) decrease in the proportion of GFP-positive leukemic cells in mice treated with CX-5461 or Dinaciclib monotherapy. A marked reduction (<95%) was observed with combination therapy, indicating decreased tumour burden across multiple compartments.

### On-Target Transcriptional Disruption and Rapid Tumour Clearance in In Vivo AML Models

To confirm on-target activity and investigate the mechanistic basis for tumour reduction in response to CX-5461 and Dinaciclib, we analysed the AML-ENL Nras-driven AML model at 3 and 10 hours post-treatment. At the 3-hour timepoint, Dinaciclib- and combination-treated mice exhibited a modest reduction in spleen size (Figure S5G), coinciding with a significant increase in apoptotic cells (Figure 5G). Concomitantly, CX-5461- and combination-treated bone marrow cells displayed a mild reduction in pre-rRNA levels (Figure 5H), while spleen cells from Dinaciclib- and combination-treated groups also showed decreased pre-rRNA expression (Figure S5H), likely reflecting a secondary effect of rapid cell death. Importantly, consistent with *in vitro* findings, phosphorylation of RPB1 at Ser2 was reduced in Dinaciclib- and combination-treated bone marrow cells, indicating effective CDK9 inhibition. Myc levels were ∼ 2X reduced in CX-5461 treated animals and variable in mice treated with combination therapy (Figure 5J).

By 10 hours post-treatment, a pronounced reduction in spleen weight was evident in Dinaciclib- and combination-treated mice (Figure 5K), accompanied by a marked decrease in bone marrow cellularity, most prominent in the combination group (Figure 5L). Circulating GFP⁺ tumour cells were significantly reduced across all treatment arms (Figure 5M), and the proportion of GFP⁺ AML cells in both bone marrow and spleen was significantly lower in Dinaciclib- and combination-treated mice (Figure 5N, Figure S5I), indicating rapid and robust tumour clearance from haematopoietic tissues. Interestingly, while CX-5461-treated mice exhibited increased levels of early and late apoptotic cells at 10 hours, this was not observed in the Dinaciclib or combination groups. Given the prominent apoptotic response already evident at 3 hours (Figure 5O), this likely reflects efficient early tumour cell death and clearance in these groups, whereas CX-5461 alone elicits a more delayed apoptotic response.

Cell cycle analysis at 10 hours further revealed significant G1-phase accumulation in total bone marrow cells, particularly in Dinaciclib- and combination-treated mice (Figure 5P). Notably, this reflects changes in the bulk cell population, in which GFP⁺ tumour cells represented less than 10% in Dinaciclib-treated animals, again consistent with early and effective tumour elimination.

Taken together, these in vivo results confirm that CX-5461 and Dinaciclib engage their respective molecular targets, suppressing RNA Polymerase I and II transcriptional outputs as anticipated. The pharmacodynamic effects are consistent with those observed in vitro and support a mechanism of synergy involving convergent transcriptional stress and rapid apoptotic clearance. The distinct kinetics of response, early cytotoxicity with Dinaciclib and delayed apoptosis with CX-5461, highlight the potential of this combination approach for effective tumour debulking in aggressive haematological malignancies.

### Therapeutic Synergy of CX-5461 and Dinaciclib in Primary Human AML

Mononuclear cells from three independent AML patients were cultured in methylcellulose-based semi-solid media and treated with either vehicle, CX-5461 (25nM), Dinaciclib (15nM), or the combination. Despite inter-patient variability in baseline colony-forming capacity, likely attributable to differences in mutational profiles (Figure S5J-L), a consistent and significant reduction in colony numbers was observed across all three samples following combination treatment. While CX-5461 and Dinaciclib monotherapies each reduced colony formation by approximately 50%, their combination led to >90% decrease in colony numbers relative to the vehicle control (Figure 5Q).

Taken together these findings highlight the synergistic effect of co-targeting RNA Pol I and CDK9 in primary human AML and support the translational potential of this combinatorial approach for treating genetically heterogeneous AML.

## Discussion

Therapeutic resistance remains a major obstacle in the treatment of aggressive haematological malignancies such as AML, where relapse rates remain high and curative options limited, particularly for patients with adverse mutational profiles including TP53 and FLT3-ITD. Our previous work demonstrated that targeting RiBi through selective inhibition of Pol I using CX-5461 monotherapy prolongs survival in AML models (Hein et al. 2017). However, durable responses were limited either by delayed progression or the emergence of acquired resistance, necessitating rational combination strategies to enhance efficacy and forestall relapse.

Here, we present a compelling preclinical case for a novel combinatorial approach that targets dual vulnerabilities, nucleolar stress and transcriptional dependency, by co-inhibiting Pol I and CDK9 using CX-5461 and Dinaciclib, respectively. Using an unbiased high-throughput screen, we identified Dinaciclib and Flavopiridol, clinically advanced CDK inhibitors, as potent enhancers of nucleolar surveillance pathway (NSP) activation. Both drugs synergized with CX-5461 in vitro and in vivo, significantly improving survival in two independent, genetically distinct murine AML models. This was paralleled by reduced tumour burden and suppression of colony-forming capacity in primary human AML samples, including those harboring mutations associated with therapy resistance, such as FLT3-ITD.

Importantly, we demonstrate that the synergistic anti-leukemic effect of CX-5461 and Dinaciclib is underpinned by their complementary mechanisms of action. CX-5461 elicits a cytostatic response through inhibition of rRNA synthesis and activation of the NSP, leading to p53 stabilization and induction of cell cycle arrest or senescence. In contrast, Dinaciclib induces rapid apoptosis by inhibiting CDK9-mediated transcriptional elongation, thereby suppressing the expression of short-lived pro-survival genes such as *MCL1*. Co-treatment with both agents results in both acute tumour cell killing and long-term proliferative suppression, an effect likely responsible for the observed delay in tumour regrowth and therapeutic resistance. These effects were not only additive at the molecular level (e.g., dual p53 stabilization, repression of MCL1), but also translated into consistent synergy across a diverse panel of AML models, regardless of driver mutation or p53 status.

Our study also reinforces the central role of the NSP as a determinant of therapeutic response. The abrogation of p53 induction in NSP-deficient settings (e.g., Mdm2^C305F^ MEFs, RPL5/11 knockdown) confirms that NSP integrity is essential for the full activity of both CX-5461 and Dinaciclib. Nonetheless, the retention of synergy in TP53-null AML cells indicates the involvement of p53-independent mechanisms, such as non-canonical stress responses. P53-independent activities have previously been reported for both CX-5461^31^ and Dinaciclib^29^, and warrant further mechanistic investigation. Our findings suggest that transcriptional or metabolic stress pathways may provide an alternative axis for synergy in p53-deficient settings, broadening the potential utility of this combination.

We also observed a modest additive effect of Dinaciclib on Pol I transcription inhibition when combined with CX-5461, despite Dinaciclib being classically described as a CDK9 inhibitor. While CDK9 is not canonically linked to Pol I activity, Dinaciclib also targets CDK1, CDK2, and CDK5, which may indirectly affect rDNA transcription. Notably, CDK1 has been shown to regulate SL-1, a core transcription factor required for Pol I transcription initiation^16^, and high concentrations of CDK inhibitors have been reported to induce nucleolar stress and disrupt Pol I regulators such as RRN3^27^. These potential cross-pathway effects merit further exploration to fully elucidate Dinaciclib’s contribution to nucleolar dysfunction.

The observation that CX-5461 also functions as a topoisomerase II poison and can induce DNA damage is a recognised limitation for clinical translation^11–13^. However, our data show that the synergistic combination with Dinaciclib permits the use of lower CX-5461 doses, thereby significantly reducing DNA damage without compromising therapeutic efficacy. The observed DNA damage-independent synergy provides further rationale for investigating combinations involving CDK9-selective inhibitors and next-generation, non-genotoxic Pol I inhibitors such as PMR-116^32^.

Taken together, our findings position CX-5461 and Dinaciclib as a rational and tractable drug combination for the treatment of genetically diverse AML. This approach targets tumour-specific vulnerabilities, including nucleolar stress sensitivity, transcriptional addiction, and apoptotic evasion, while also mitigating toxicity associated with high-dose monotherapies. Unlike our prior studies using MTD CX-5461^26^ dosing, our work demonstrates that combining lower, sub-cytotoxic concentrations of CX-5461 and Dinaciclib yields superior efficacy with reduced genotoxic burden. Importantly, this strategy not only enhances anti-leukaemic activity but also minimizes adverse effects such as DNA damage, supporting its potential use in older or medically frail patients who are typically ineligible for intensive chemotherapy.

While residual disease was still observed at late timepoints, particularly in sanctuary sites such as the cranial region, the combination regimen substantially delayed disease progression and improved overall survival compared to monotherapies. These findings reinforce the translational utility of the approach, particularly in settings where standard MTD regimens are poorly tolerated.

In conclusion, we provide robust preclinical evidence that co-targeting RNA Polymerase I and CDK9-mediated transcriptional elongation represents a highly effective and tolerable therapeutic strategy for high-risk AML. By enabling disease control at lower drug doses and across genetically heterogeneous disease, this combination holds promise for broader clinical application, including in patient populations with limited treatment options, and warrants further clinical development, including trials of next-generation, non-genotoxic Pol I inhibitors. Dinaciclib, a clinically advanced CDK inhibitor currently in Phase III trials for refractory CLL, has demonstrated activity in multiple cancer types, including AML, and thus offers an attractive, well-characterised partner for rational combination strategies such as the one described here.

## Material and methods

### Cell lines

A549 (ATCC, CCL-185), Mv4;11 (ATCC, CRL-9591), NOMO-1 (DSMZ, ACC 542), MOLM1-3 (DSMZ, ACC 554), SHI-1 (DSMZ, ACC 645), KG-1 (DSMZ, ACC 14), OCI-AML3 (DSMZ, ACC 582) were grown according to supplier instructions. MDM2 WT and MDM2 C305F were establish from Mdm2^tm1.1Ypz^ mice^28^ (Kindly provided by Prof Yanping Zhang (University of North Carolina, Chapel Hill, USA) and cultured as described^33^ under ANU AEC-approved animal ethics protocol (A2018/56). Cells were maintained bellow 80% confluency (adherent) or 2×10^6^ cells/ml (suspension) in a humidified 5% CO_2_ incubator at 37°C. Cells were seeded 24 hours prior to treatment to ensure exponentially growing phase at the time of experiment.

### High-throughput compound screening

Screen was performed has previously decribed^28^ with some adaptations. At the commencement of screening (Day 1), A549 cells were reverse transfected with 10 nM siGENOME human siRPS19 SMARTpool siRNA (GE Dharmacon, M-003771) into 384 well black-walled optical microplates (Corning Costar 3571) and incubated for 24 hours (37°C, 5% CO_2_, humidified). On Day 2, medium (DMEM/F12, 2mM L-glutamine L-glutamine,10% FBS) was replaced and DMSO or compounds added to wells to final concentration of 5uM, 1uM, 200nM and 40nM. The compound library (4169 FDA-approved compounds) was obtained from the Walter and Eliza Hall Institute Drug Discovery Centre (via Compounds Australia, Griffith University, Australia). Cells were incubated with compounds for 48h. On Day 4 (48 hours post-drug treatment), plates were fixed with 4% Paraformaldehyde (Proscitech C0042B), stained with 0.5 µg/mL 4′,6-diamidino-2-phenylindole (DAPI, D9542, Sigma Aldrich) and permeabilised with 0.5% Triton X-100 (Sigma Aldrich TTX-100). Cells were incubated with anti-p53 and anti-mouse A594 (See table 3) before being imaged on the Cellomics ArrayScan VTi (Thermo) using the following parameters: image acquisition was carried out using the 20X objective in standard mode (1024×1024; 2×2 binning), capturing 25 fields/well at a fixed exposure time. Images were processed using a Cellomics Target Activation Bioapplication developed to identify DAPI-stained nuclei (as a surrogate to count cell number). The nuclear mask was then used to quantify the average nuclear intensity of p53 staining per cell, which was then translated into an average/well as a screening readout and normalised to the DMSO only control. The Mahalanobis distance ^34^ was used as a metric to score compounds according to their therapeutic potential, i.e. increased p53 expression and decreased cell number over the four different concentrations.

### siRNA transfection

A549 cells were reverse transfected with 5nM NT (L-013549), RPS19 (L-003771), RPL5 (L-013611) or RPL11 (L-013703) SMARTpool siRNA and DharmaFECT1 (T-2001) as per manufacturer instructions (Dharmacon). After 24 hours of transfection, media was replaced with A549 growth media containing either vehicle, single drug or drug combination. Cells were incubated for further 48 hours prior to harvest for Protein or RNA expression analysis.

### Inhibitors

Flavopiridol (Selleckchem S1230) and Dinaciclib (Selleckchem S2768) were dissolved in 100% DMSO and CX-5461 was dissolved in 50mM NaH2PO4 pH4.5, and stored at −20oC until experimental use. Inhibitor stocks were diluted to the required concentrations in corresponding vehicles: 50 M NaH2PO4 or 0.1% DMSO for single drug assays, or 50 μM NaH2PO4 + 0.1% DMSO for combination assays.

### Target inhibitory concentration (TIC50) determination

Cell lines were seeded 24 prior to drug treatment and treated with a range of drug concentrations for 3 hrs. For Pol I transcription inhibition IC50 determination, RNA expression was analysed as described in “RNA expression analysis by qPCR” using primers for 5’ETS, cFOS and B2M (see Table 2). For CDK inhibition, protein extracts were prepared and Phospho-RPB1 (Ser2) and p53 expression (see Table 3) were analysed as described in “Protein expression analysis by WB”. The expression changes were further normalised to the vehicle and the dose response curve obtained with GraphPad Prism 10 using non-linear regression ([inhibitor] versus response – variable slope (four parameters)) and the target IC50 (TIC50) calculated.

**Table 2.**
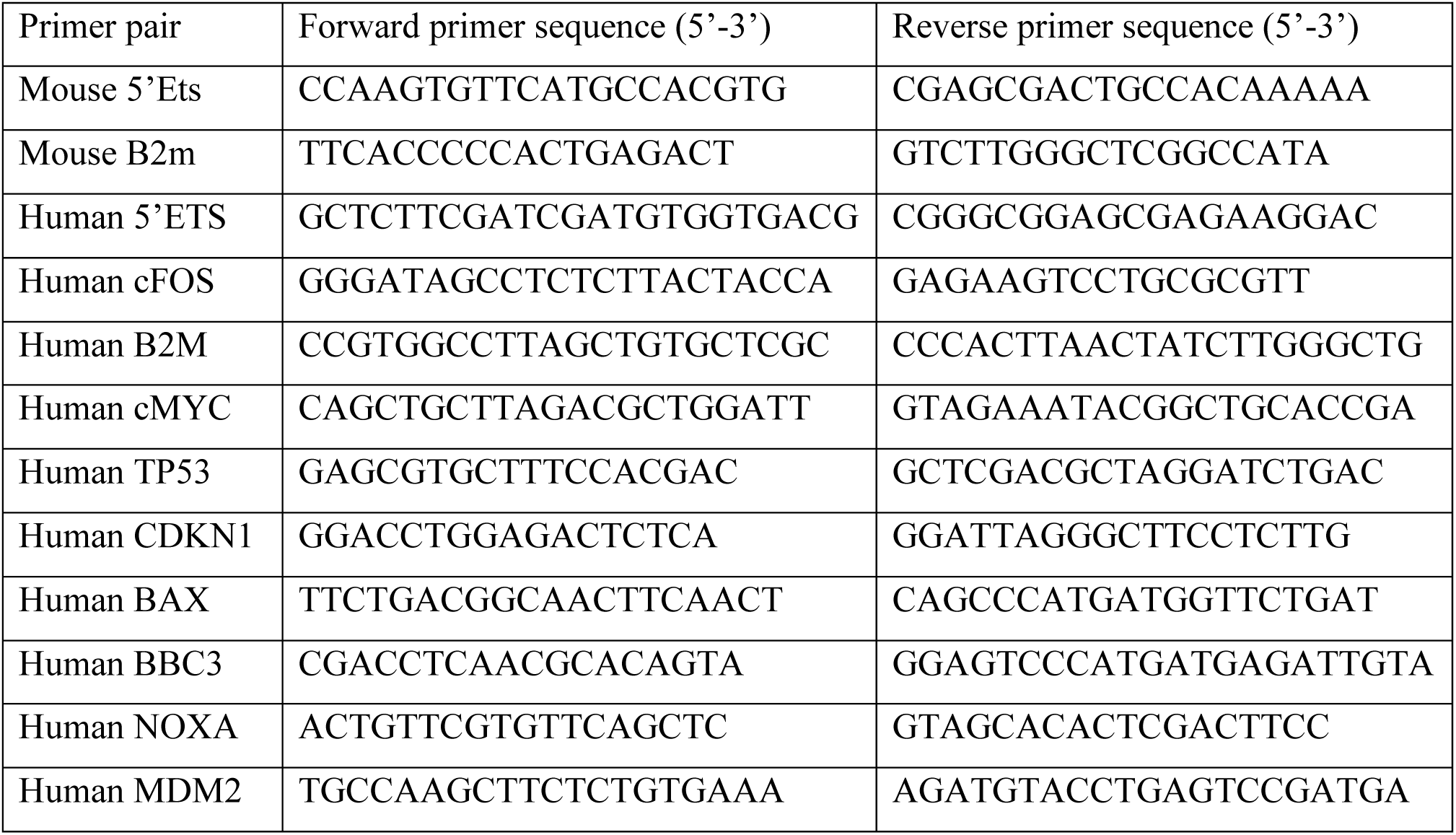
Primers.

### Growth inhibitory concentration (GIC50) determination

Cell lines were treated with a range of drug concentrations and grown until vehicle treated culture reached 100% confluency. Images were collected every 6 hours using Incucyte Zoom (Essen BioScience 2016A) life cell imaging system. Images were analysed with Incucyte Control software (Essen BioScience) to determine percentage confluency at different time points. Confluency values at the time point vehicle-treated cell reach confluency were plotted and the dose response curve obtained with GraphPad Prism 10 using non-linear regression ([inhibitor] versus response – variable slope (four parameters)) and the growth IC50 (IC50) calculated.

### Cell proliferation assays

A549 cells treated with vehicle, CX-5461, Dinaciclib/Flavopiridol or combination were imaged every 6 hours and confluency calculated as described in “GIC50 determination”. Confluency values at the time point vehicle-treated cell reach confluency were used for analysis. Mv4;11 cells treated with vehicle, CX-5461, Dinaciclib or combination were counted every 24h from the time treatment started using Z series CoulterCounter (Beckman Coulter). Cell counts/confluency values were normalized to the time when treatment was started (T=0h).

### RNA expression analysis by quantitative polymerase chain reaction (qPCR)

After 3 hours treatment with vehicle, single drug or drug combinations, total RNA was extracted using a Nucleospin RNA kit (Macherey-Nagel 740955), cDNA was prepared using SuperScript IV Reverse Transcriptase (Invitrogen) and qPCR performed on the StepOne Plus Real-Time PCR System (Applied Biosystems) using a SYBRGreen Fast Master Mix (Applied Biosystems) and target specific primers (see Table 2). For each replicate, RNA expression was normalised the housekeeping gene B2M via the 2^−(ΔCt)^ method. Expression changes were further normalised to vehicle (treatment/vehicle).

### Protein expression analysis by immunoblotting

After 3 hours treatment with a single drug or drug combinations, whole cell protein extracts were prepared in 20mM HEPES, 0.5mM EDTA, 2% SDS, 0.1M DTT buffer. Proteins were separated by SDS-polyacrylamide gel electrophoresis using 4-20% Mini-PROTEAN TGX Precast Protein gels (Bio-Rad 4561093) and transferred to Immobilon-P PVDF membrane (Millipore, IPVH00010) using Trans-Blot Turbo Transfer System (BioRad) or using iBlot Transfer Stack with PVDF membrane (IB24001) using iBlot2 Gel Transfer Device (Invitrogen IB21001). Membranes were probed with target-specific antibodies (see Table 3) and visualised by using Clarity Western ECL Substrate (BioRad 1705061) in the ChemiDoc System (BioRad). Integrated volume for antibody-specific bands were calculated with Image Lab software (version 6.0.1, 2017) and values were normalized to housekeeping protein Beta-actin (ACTB) or Vinculin (VCL) (see Table 3).

**Table 3.** Antibodies.

| **Antibody ID** | **Reactivity** | **Cat number** | **Supplier** |
| --- | --- | --- | --- |
| RPB1 phospho-CTD (S2) | human, mouse | 13499S | Cell Signalling |
| p53 | human | sc-126 | Santa Cruz Biotech |
| P21 | human | 2947S | Cell Signalling |
| MCL1 | human, mouse | ab32087 | Abcam |
| Cleaved CASPASE 3 | human, mouse | 9661 | Cell Signalling Technologies |
| PARP | human, mouse | 9541 | Cell Signalling Technologies |
| FIBRILLARIN | human, mouse | ab5821 | Abcam |
| NUCLEOLIN | human, mouse | ab136649 | Abcam |
| phospho-γH2AX | human, mouse | 05-636 | Merk |
| BETA ACTIN HRP (ACTB) | human, mouse | sc-47778 | Santa Cruz Biotech |
| VINCULIN-HRP (VCL) | Human, mouse | 18799 | Cell Signalling Technologies |
| BrdU | N/A | 347580 | BD Biosciences |
| IgG-Alexa Fluor 488 | rabbit | A11008 | Life Technologies |
| IgG- Alexa Fluor 488 | mouse | A11001 | Life Technologies |
| IgG- Alexa Fluor 647 | mouse | A21235 | Life Technologies |

### Protein expression analysis by immunofluorescence (IF)

After the required incubation with the different single drug or drug combinations, cells were fixed with 4% Paraformaldehyde (PFA; Electron Microscopy Sciences 15714S), permeabilized with 0.5% Triton X100, blocked with 5% bovine serum albumin (BSA) in phosphate buffer saline (PBS). Primary antibodies were incubated either 1h at RT or 16h at 4°C and secondary antibodies were incubated for 1h at RT (except for gH2AX staining where secondary was incubated at 37oC) and stained with 5μg/ml DAPI (Sigma D-9542). A549 cells were cultured, treated and stained on glass coverslip or Phenoplate 96 well microplates (Revvity, 6055302). Mv4;11 were cultured and treated in multi-well plates and cytospun onto PLL coated glass slides (Sigma-Aldrich P0425) or Poly-L-lysine (PLL)-coated Phenoplate 96 well microplates (Revvity, 6055302). Slides were mounted using Fluorescence Mounting Media (Agilent S3023) and imaged using LSM800 confocal microscope (Zeiss) with a 63X (1.4) lens. Cells in 96 well plates were kept in PBS plate were sealed and imaged in Opera Phenix High-Content Screening System (PerkinElmer) with a 40X lens.

### Viability analysis

Cells were treated with vehicle, 0.1 to 10X IC50 of CX-5461 and Dinaciclib alone or in combination for 72h. To assess cell viability, cells were stained with 1μg/ml Propidium Iodine (PI; Sigma-Aldrich 287075), data was acquired with FACSVerse (BD Life Sciences) and analysed using FlowJo^TM^ Software. Synergy was evaluated using Chou-Talalay CI values, calculated with the CalcuSyn software 2.1 (Biosoft) as previously described^35^. The lowest CI value obtained corresponding to Fa value greater than 0.5 was used to assess the degree of synergy (CI<1), additivity (CI=1) or antagonism (CI>1).

### Apoptosis analysis

Cells were stained at 3, 6 and 24h post-treatment with vehicle, CX-5461, Dinaciclib or in combination using APC Annexin V Apoptosis Detection Kit (BioLegend 640920). Data was acquired with BD LSR II flow cytometer (BD Life Sciences) and analysed using FlowJo^TM^ v10.8 Software. Synergy was assessed using Bliss independent model and synergy defined by Excess over Bliss Combination Index (CI) > 0.

### Cell cycle analysis

Cells were analysed at 12 and 24h post-treatment with vehicle, CX-5461, Dinaciclib or in combination. Thirty minutes prior to harvest, cells were pulse-labelled with 10uM BrDU. After harvesting, cells were fixed with ice cold 80% ethanol, treated with 2N HCl/ 0.5% (v/v) Triton X100 and neutralized with 0.1M Na_2_B_4_O_7_.10H2. BrdU incorporation was probed with anti-BrDU antibody and anti-IgG-A488 and DNA was stained with 10ug/ml Propidium Iodine. Data was acquired with APF-LSR II flow cytometer (BD Life Sciences) and analysed using FlowJo^TM^ v10.8 Software.

### Synergy analysis

Except when stated otherwise (viability and apoptosis analysis), synergy scores were calculated using SynergyFinder V3 (https://synergyfinder.fimm.fi/^36^) using Bliss independence model. Synergy was defined by Bliss score > 10.

### *In vitro* transcription assays

Non-specific transcription assays (NSA) were performed as described^37^.

Promoter driven run-off transcription assays (PSA) were performed as described^38^. Briefly, 200ng of linear 440 bp DNA fragment containing the human rRNA gene promoter were used as template. Reactions were supplemented with 5μl of HeLa Nuclear extract (NE) and vehicle (DMSO), Dinaciclib or Flavopiridol. Transcription was initiated with 500 μM UTP, GTP and ATP, 25 μM CTP and 2.5 μCi [α32P]CTP (3000 Ci/mmole), and 2 U RNasin, 0.1 mg/ml α-amanitin, 10 mM creatine phosphate. After 45 min at 30^0^C, 10 U RNase-free DNase I (Roche) was added and incubated at 37^0^C for 5 min. Reactions were terminated at 37^0^C for 5 min with 200 μl of 20 mM EDTA, 200 mM NaCl, 1% (w/v) SDS, 0.25 μg/μl tRNA and 20 mg/ml Proteinase K. Nucleic acids were phenol-chloroform extracted, ethanol precipitated, dissolved in formamide loading buffer and analysed on denaturing (8 M urea) 11% polyacrylamide gels. Drugs (or vehicle) were added prior of addition of NTP’s.

The amount of RNA produced by in vitro transcription reactions was quantified using a phosphorimager, FLA-7000 (Fuji) and Aida software.

### MLL/ENL Nras and MLL-AF9 AML combinatorial treatment

For survival experiments, 8-10 weeks old female C57BL/6 mice were transplanted with 5e^5^ MLL/ENL Nras (p53 WT) or MLL-AF9 AML cells via tail vein injection. Seven days post-transplant, the mice were randomized into four groups and dosed continuously with either CX-5461 vehicle (CX-5461 - 50mM Phosphate buffer in PBS (pH 4.5)) or 30mg/kg CX-5461 (3 days/week) by oral gavage; CDK inhibitor vehicle (20% W/V 2-Hydroxypropyl-β-cyclodextrin in PBS (Sigma-Aldrich C0926), 20mg/kg Dinaciclib or 7.5mg/kg Flavopiridol by interperitoneally injection (2 day/week) or a combination of both CX-5461 and CDK9 drugs until an ethical endpoint was reached. Engraftment and disease progression were monitored once a week (starting at day of treatment) by bioluminescence imaging using the IVIS Xenogen after injection of mice with luciferin (200µl per mouse of a 1mg/ml stock solution). Mice were weight at least three times a week.

For short term therapy experiments, mice were transplanted as described above and treated when disease was established with a single dose of vehicles, CX-5461, Dinaciclib or combination of both. Peripheral blood was collected 10h post-treatment, whole blood cell counts were performed with ADVIA 120 Hematology System (Siemens) and circulating tumour burden was evaluated by flow cytometry. Bone marrow and spleen tissues were harvest at 3 and 10 hours post-treatment for RNA and protein expression analysis, tumour burden, apoptosis and cell cycle analysis. Three technical replicates where performed for each patient sample. Experiments were conducted under institutionally approved animal ethics protocols (ANU: A2015/12 & A2018/48; Peter MacCallum Cancer Centre: E462).

### Primary human AML colony formation assay

Mutation analysis was performed in de-identified AML patient biopsy samples to identify AML driver mutations as described previously^39,40^. Colony formation capacity of primary human AML cells was analysed in methylcellulose semi-solid media (Stem Cell Technologies M4435) as described previously^5^. Bone marrow cells (10^5^) were plated and treated with vehicles, 25nM CX-5461, 15nM Dinaciclib or combination of both drugs. Patient consent was obtained under UniSA human research ethics number ETHLR16.141.

## Acknowledgements

We would like to thank the staff at the Australian Phenomics Facility, Dr. Harpreet Vohra and Mr. Michael Devoy at the Cytometry, Histology and Spatial Multiomics Facility, Dr. Jinshu He at the ANU Centre for Therapeutic Discovery (ACTD) and Dr Daryl Webb at the Centre for Advanced Microscopy at the Australian National University. We would also like to thank staff at the Victorian Centre for Functional Genomics at the Peter MacCallum Cancer Centre, Compounds Australia and the WEHI Drug Discovery Centre (now known as the National Drug Discovery Centre; NDDC). The ACTD, VCFG, Compounds Australia and NDDC receive operational funding from the Australian Government’s National Collaborative Research Infrastructure Strategy (NCRIS) program through Therapeutic Innovation Australia, while the ACTD and VCFG also receive funding from Phenomics Australia. SSM was supported by the Brazilian Ministry of Education via the Coordination for the Improvement of Higher Education Personnel (CAPES) foundation. This work was supported by funding from the National Health and Medical Research Council (NHMRC) of Australia (Project Grants 1158732, Ideas Grant 2002741 and Senior Research/Investigator Fellowships to RDH (1116999, 2009504).

## Authors contribution

NH, LF, KP, KMH, AJG, RDH and RF conceived, designed and supervised the study. RF wrote the manuscript; all other authors have reviewed and edited the manuscript. Screen was designed by RDH and AJG, executed by AJG, SJA, KMG, PM and KJS, and analysed by AJG, KJS, ME and AC; Preclinical studies were conducted and analysed by NH, LNV, SSM, JS, KMH and RF; In vitro experiments were performed by NH, PP, JS, JP, PS, JF, DT, KP and RF.

## Notes

### Competing Interest Statement

The authors have declared no competing interest.

