## Supplemental figures and legends for "Novel RNA Polymerase I and Cyclin Dependent Kinase combination therapy for the treatment of aggressive Acute Myeloid Leukemia"

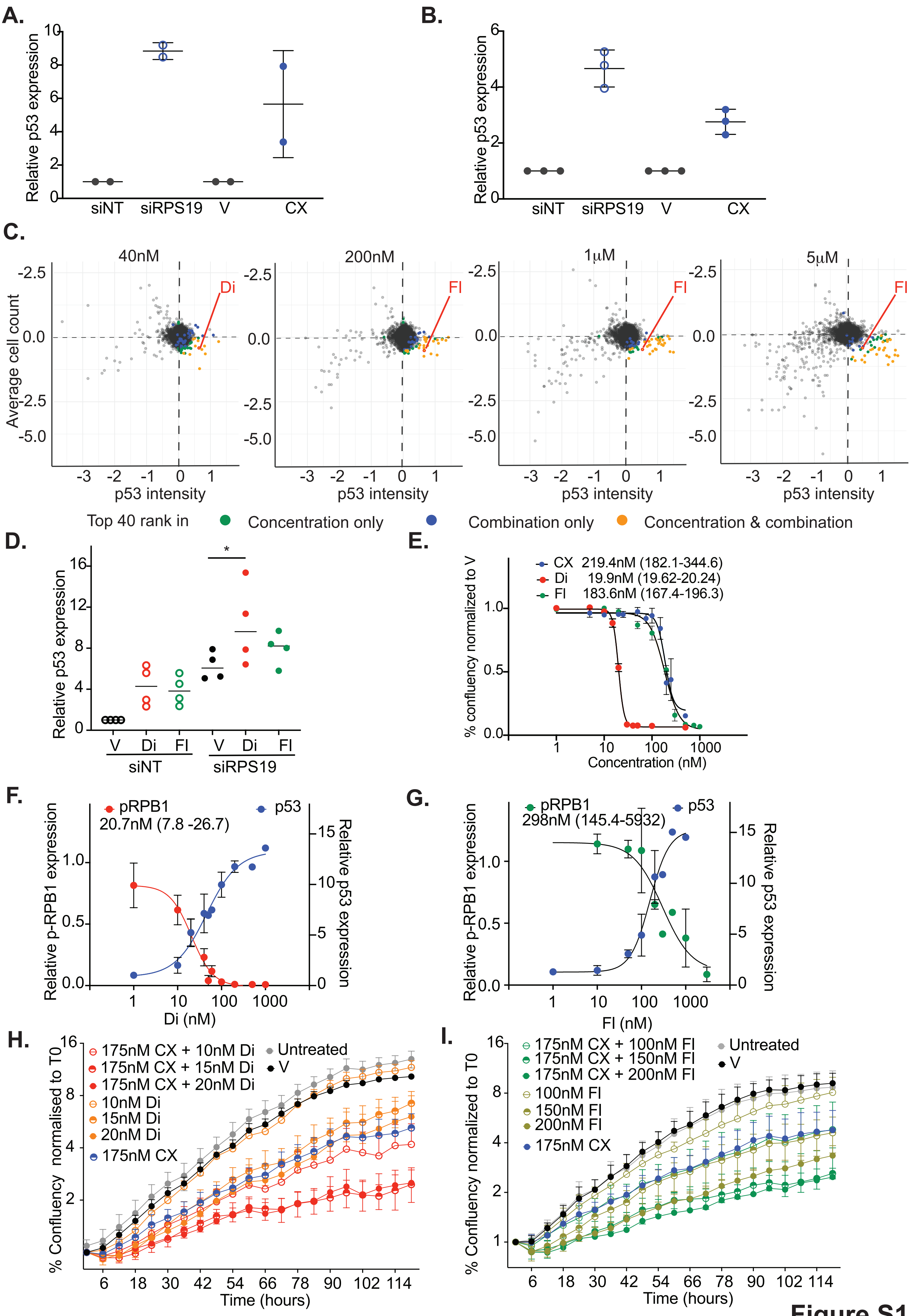
A.
Relative p53 expression
siNT
siRPS19
V
CX
B.
Relative p53 expression
siNT
siRPS19
V
CX
C.
40nM
200nM
1μM
5μM
Di
FI
Average cell count
p53 intensity
Top 40 rank in
Concentration only
Combination only
Concentration & combination
D.
Relative p53 expression
V
Di
FI
siNT
siRPS19
E.
CX 219.4nM (182.1-344.6)
Di 19.9nM (19.62-20.24)
FI 183.6nM (167.4-196.3)
% confluency normalized to V
Concentration (nM)
F.
pRPB1
20.7nM (7.8 -26.7)
p53
Relative p-RPB1 expression
Relative p53 expression
Di (nM)
G.
pRPB1
298nM (145.4-5932)
p53
Relative p-RPB1 expression
Relative p53 expression
FI (nM)
H.
175nM CX + 10nM Di
175nM CX + 15nM Di
175nM CX + 20nM Di
10nM Di
15nM Di
20nM Di
175nM CX
Untreated
V
% Confluency normalised to T0
Time (hours)
I.
175nM CX + 100nM FI
175nM CX + 150nM FI
175nM CX + 200nM FI
100nM FI
150nM FI
200nM FI
175nM CX
Untreated
V
% Confluency normalized to T0
Time (hours)

**Figure S1**

**Figure S1 A-B.** Quantification of immunoblotting (A) and immunofluorescence (B) shown in figures 1A and 1B, respectively. **C**. Distribution of Mahalanobis distance of each library compound across 40nM, 200nM, 1μM and 5μM drug concentrations. Top 40 ranked drugs identified only in a single concentration, only in the combined data from all concentrations tested or in both single and combine data depicted in green, blue and orange, respectively. Dinaciclib was identified at 40nM concentration as well as combined concentrations. Flavopiridol was identified at 200nM, 1μM and 5μM concentrations as well as in combined concentrations. **D.** Quantification of p53 protein abundance in A549 cells transfected for 48h with siNT or siRPS19 and treated for 3h with Dinaciclib or Flavopiridol. Densitometry values were normalized to Vehicle within each biological replicate. **E-F**. Quantification of p53 and p-RBP1 protein abundance in A549 cells treated for 3h with 8 concentrations of Dinaciclib (1nM to 1μM). Densitometry values were normalized to Vehicle within each biological replicate and target IC50 was calculated. **G**. Dose response curves showing percentage confluency of A549 cells treated with 10 different concentrations of CX-5461, Dinaciclib and Flavopiridol (from 0.3nM to 10μM), normalize to vehicle used for growth IC50 calculation. Graphs represent mean +/- SD of n= 3 independent experiments. **I-J.** Percentage confluence over time, normalized to time 0, of A549 cells treated with 175nM CX and variable concentrations of Dinaciclib (10-20nM; H) and Flavopiridol (100-200nM;G). Graphs represent mean +/- SEM of 3 independent experiments.

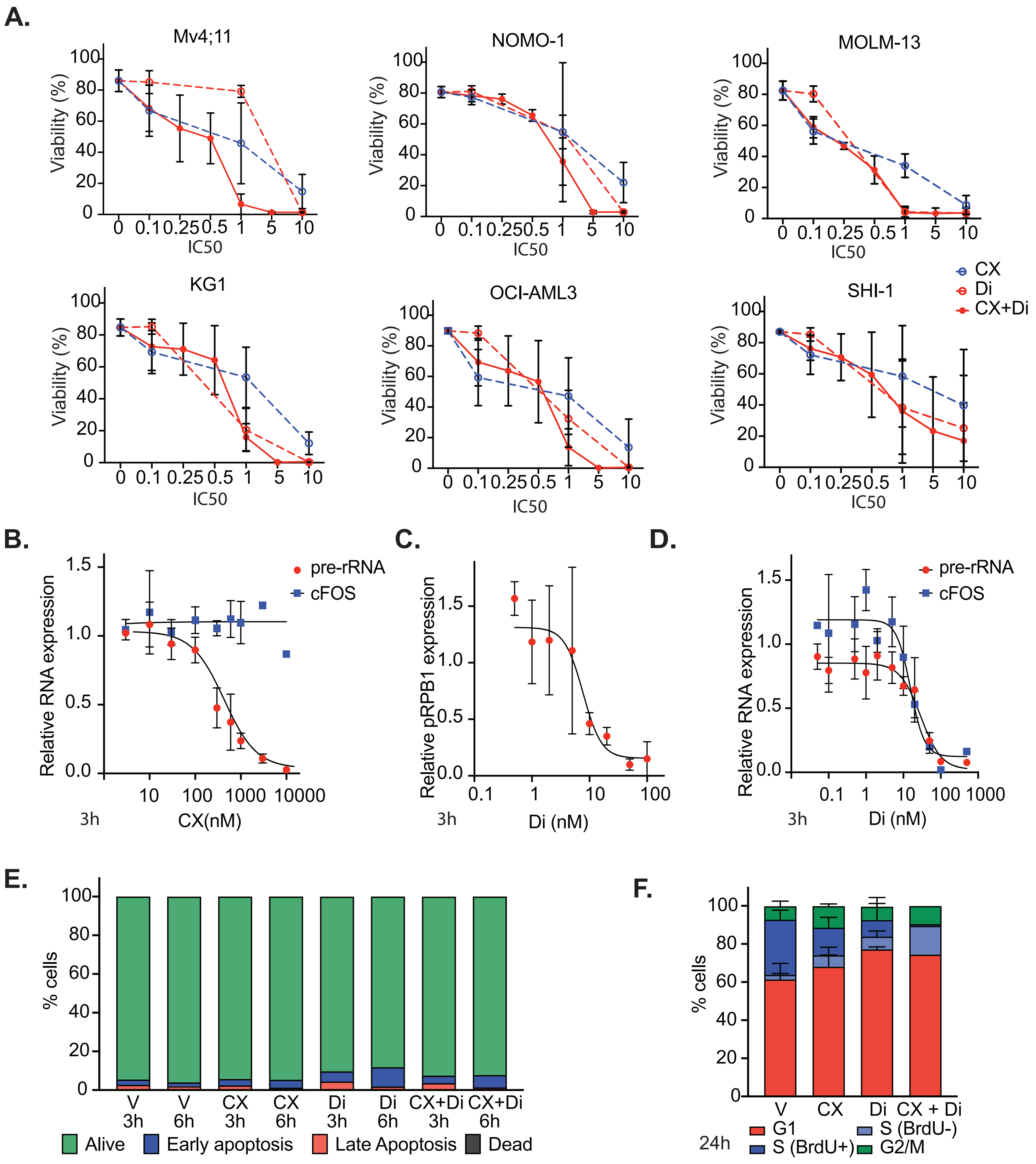
A.
Mv4;11
NOMO-1
MOLM-13
KG1
OCI-AML3
SHI-1
Viability (%)
IC50
0 0.1 0.25 0.5 1 5 10
CX
Di
CX+Di
B.
Relative RNA expression
pre-rRNA
cFOS
3h
CX(nM)
C.
Relative pRPB1 expression
3h
Di (nM)
D.
Relative RNA expression
pre-rRNA
cFOS
3h
Di (nM)
E.
% cells
V 3h
V 6h
CX 3h
CX 6h
Di 3h
Di 6h
CX+Di 3h
CX+Di 6h
Alive
Early apoptosis
Late Apoptosis
Dead
F.
% cells
V
CX
Di
CX + Di
24h
G1
S (BrdU-)
S (BrdU+)
G2/M

Figure S2

**Figure S2 A.** Dose response curves showing percentage viability in human AML cell lines treated with variable CX-5461 and Dinaciclib concentrations. Concentration used are factor of the pre-determine IC50 values (Baker et al. 2016, Hein et al. 2017). Mv4,11 & MOLM-13: 300nM CX-5461, 20nM Dinaciclib; NOMO-1: : 65nM CX-5461, 20nM Dinaciclib; KG1, OCI-AML3, SHI-1 and OCI-M2: 930nM CX-5461, 20nM Dinaciclib. Graphs represent mean +/- SD of n= 3 independent experiments. **B.** Relative pre-rRNA (Pol I transcribed gene) and cFOS (PolII transcribed gene) expression of Mv4;11 cells treated with 10 concentrations of CX-5461 for 3 hours (Ferreira et al. 2025). Dose response curve was used to calculate target IC50 for CX-5461 (300nM). **C.** Quantification of p-RPB1 protein abundance in Mv4;1 cells treated for 3h with 8 concentrations of Dinaciclib (0.3nM to 100nM). Densitometry values were normalized to vehicle within each biological replicate and target IC50 was calculated for Dinaciclib (10nM). **D.** Dose response curves showing pre-rRNA and cFOS RNA expression in Mv4;11 cells treated with 11 different concentrations of Dinaciclib and (from 0.03nM to 1µM), normalize to vehicle showing a concentration-dependent reduction in both cFOS and rRNA expression. Graphs represent mean +/- SD of n= 3 independent experiments. **E.** Apoptosis analysis of Mv4;11 treated for vehicle, 300nM CX-5461, 10nM Dinaciclib and combination of both drugs at 3 and 6 hours post-treatment showing low levels of apoptosis (n=1). **F.** Cell cycle analysis of Mv4;11 cell treated for 24 hours with vehicle, 300nM CX-5461, 10nM Dinaciclib and combination of both drugs. Sub-G1 (dead) cells were excluded from analysis. Graphs represent mean +/- SD of n= 3 independent experiments with the exception of combo treatment where 2 biological replicates show very high levels of cell death presenting analysis.

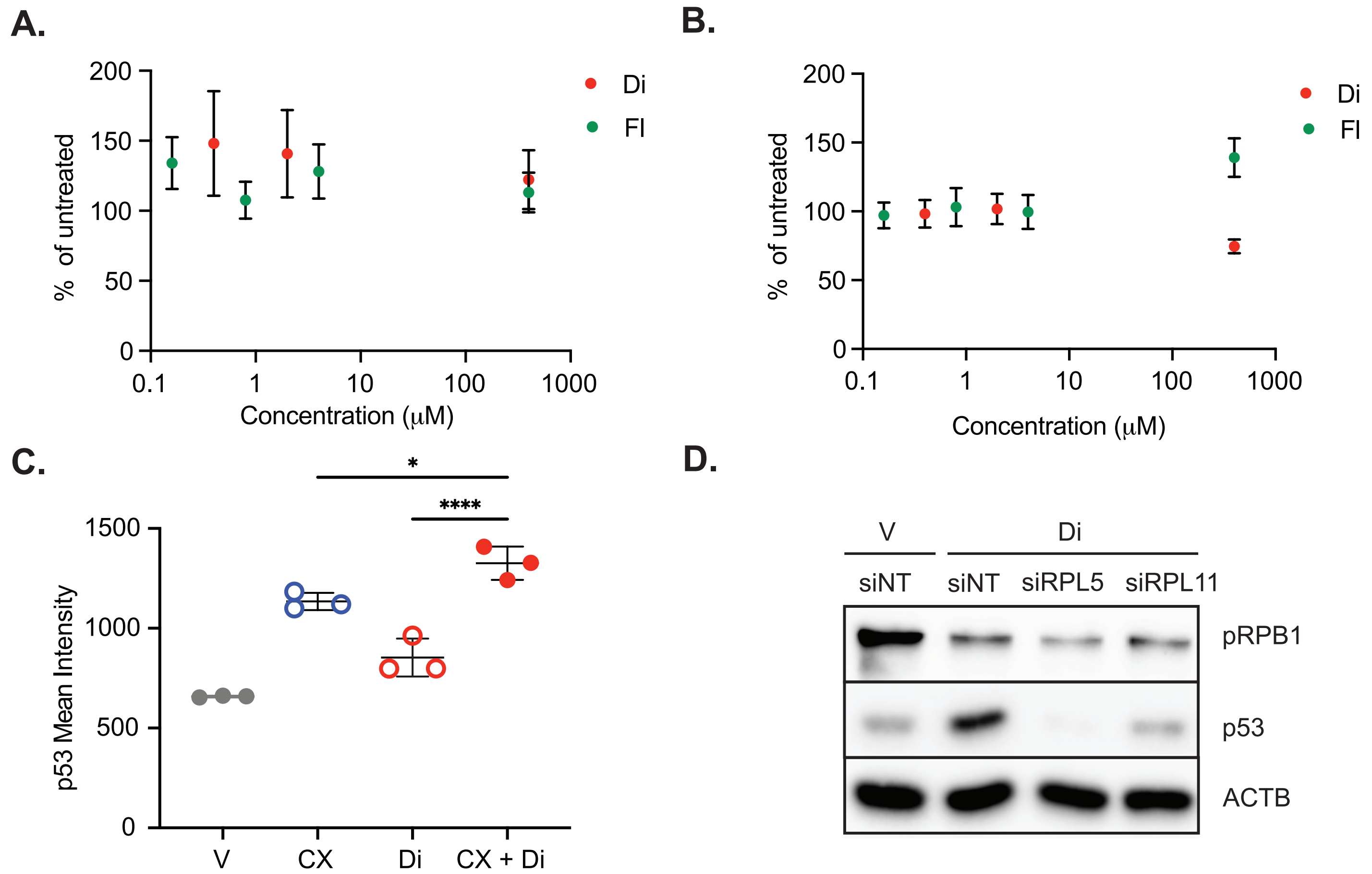
A.
% of untreated
Concentration (μM)
Di
Fl
B.
% of untreated
Concentration (μM)
Di
Fl
C.
p53 Mean Intensity
V
CX
Di
CX + Di
*
****
D.
V
Di
siNT
siNT
siRPL5
siRPL11
pRPB1
p53
ACTB

Figure S3

**Figure S3 A.** Non-specific transcription assay (NSA) and **B.** Promoter specific assay (PSA) performed using a cell-free system with HeLa nuclear extract and treatment with variable concentrations of Dinaciclib and Flavopiridol. **C.** Mean p53 intensity per well in Mv4;11 cells were treated for 3h with vehicle and TIC50 concentrations of CX-5461 (300nM), Dinaciclib (10nM) or combination of both drugs. Graph represent mean +/- SD of 3 independent experiments. **D.** Representative immunoblot of p53 protein levels protein and ACTB in A549 with and without knockdown of RPL5 or RPL11.

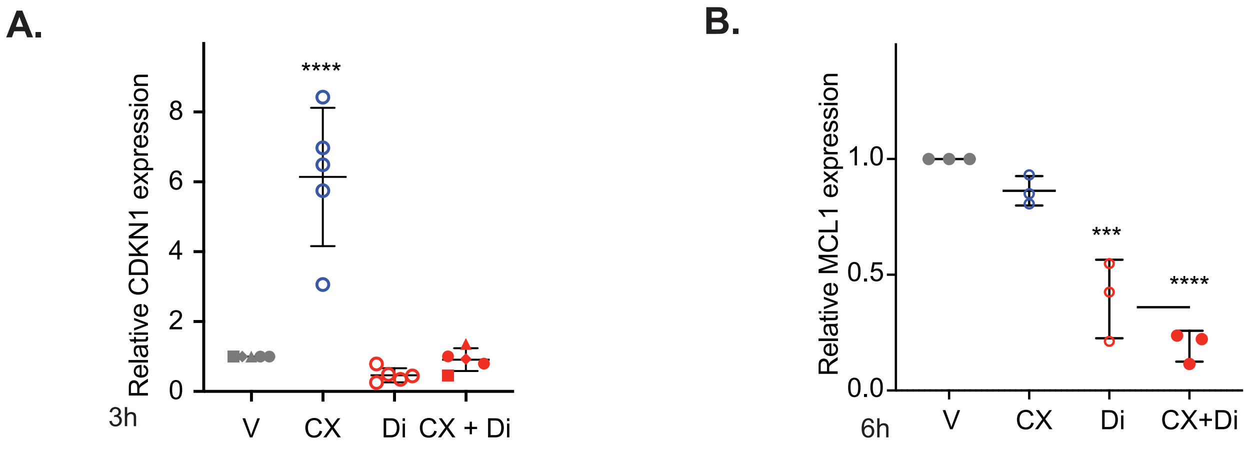
A.
Relative CDKN1 expression
0
2
4
6
8
****
3h
V
CX
Di
CX + Di
B.
Relative MCL1 expression
0.0
0.5
1.0
***
****
6h
V
CX
Di
CX+Di

**Figure S4**

**Figure S4 A.** Quantification of CDKN1 (p21) protein level, normalized to BACT and vehicle 3h post-treatment with CX-5461 (300nM), Dinaciclib (10nM) or combination of both drugs. **B.** Quantification of MCL1 protein level, normalized to BACT and vehicle 6h post-treatment with CX-5461 (300nM), Dinaciclib (10nM) or combination of both drugs.

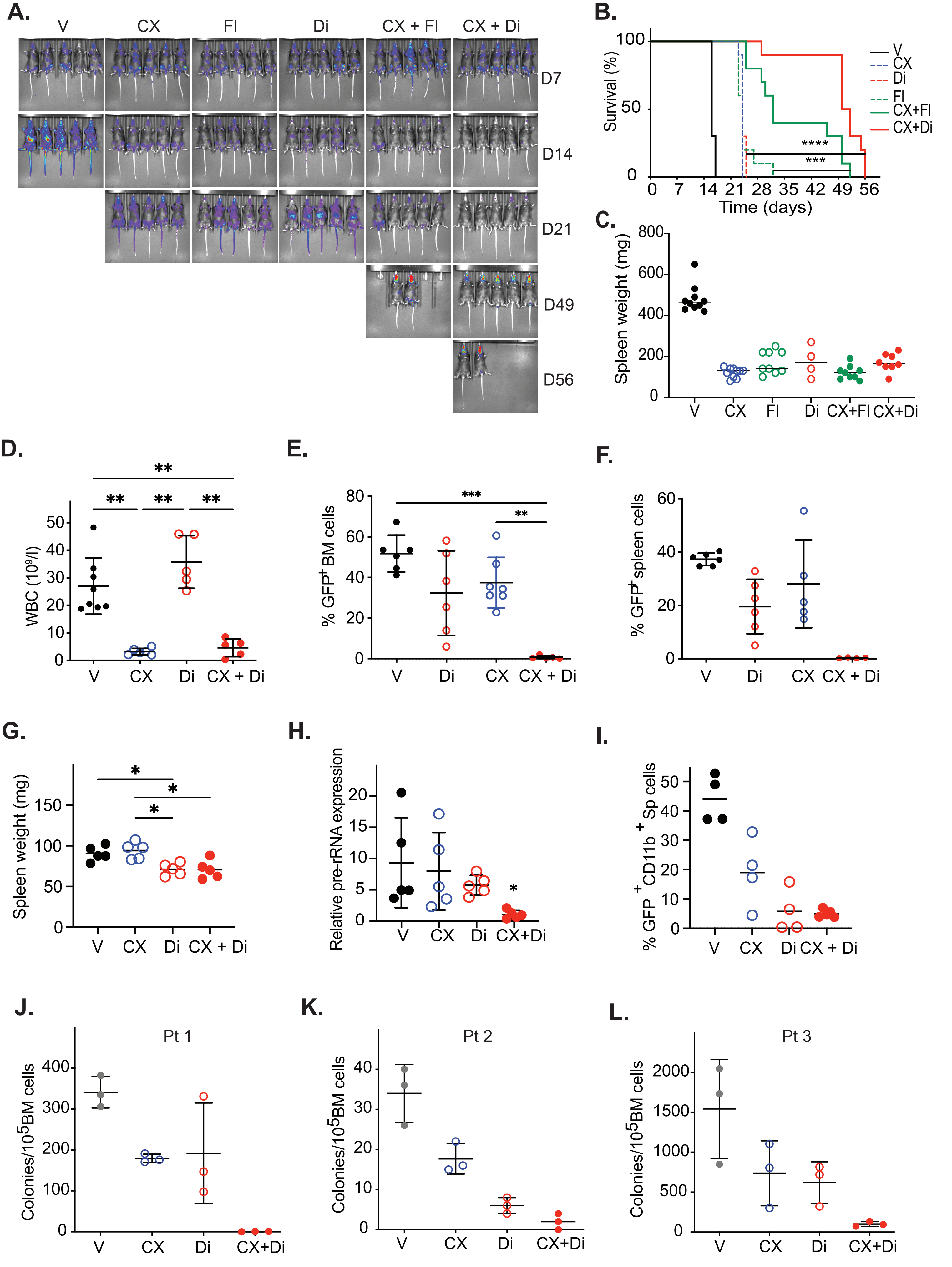

A.
V
CX
FI
Di
CX + FI
CX + Di
D7
D14
D21
D49
D56
B.
Survival (%)
Time (days)
V
CX
Di
FI
CX+FI
CX+Di
****
***
C.
Spleen weight (mg)
V
CX
FI
Di
CX+FI
CX+Di
D.
WBC (10⁹/l)
V
CX
Di
CX + Di
E.
% GFP+ BM cells
F.
% GFP+ spleen cells
G.
Spleen weight (mg)
H.
Relative pre-rRNA expression
I.
% GFP+ CD11b+ Sp cells
J.
Pt 1
Colonies/10⁵BM cells
K.
Pt 2
L.
Pt 3


**Figure S5**

**Figure S5 A.** Representative images of bioluminescence imaging used to monitor disease progression **B**. Kaplan-Meier plot showing overall survival and **C**. Spleen weight **D**. Peripheral white blood cells count **E.** Percentage tumour (GFP+) cells in bone marrow and **F.** Percentage tumour (GFP+) cells in spleen at ethical endpoint of MLL-AF9 $Nras^{G12D}$ AML model treated with vehicle, CX-5461 or Dinaciclib monotherapy or combination therapy. **G.** Spleen weights H. Relative pre-rRNA and **I.** Percentage of tumour cells (GFP+ CD11B+) in spleen 10h post-treatment of MLL/ENL Nras treated with vehicle, CX-5461 or Dinaciclib monotherapy or combination therapy. J-L. Absolute colony number for AML patient 1, 2 and 3 bone marrow cells treated with vehicle, 25nM CX-5461, 15nM Dinaciclib or combination of both drugs. All graphs display mean ± standard deviation of at least 3 biological (A-I) or technical (J-L) replicates.
